# Developmental exposure to domoic acid targets reticulospinal neurons and leads to aberrant myelination in the spinal cord

**DOI:** 10.1101/2022.08.08.503200

**Authors:** Jennifer M. Panlilio, Katherine M. Hammar, Neelakanteswar Aluru, Mark E. Hahn

## Abstract

Harmful algal blooms (HABs) produce neurotoxins that affect human health. Developmental exposure of zebrafish embryos to the HAB toxin domoic acid (DomA) causes myelin defects, loss of reticulospinal neurons, and behavioral deficits. However, it is unclear whether DomA primarily targets myelin sheaths, leading to the loss of reticulospinal neurons, or reticulospinal neurons, causing myelin defects. Here, we show that while exposure to DomA at 2 dpf did not reduce the number of oligodendrocyte precursors prior to myelination, it led to fewer myelinating oligodendrocytes that produced shorter myelin sheaths and aberrantly wrapped neuron cell bodies. DomA-exposed larvae lacked Mauthner neurons prior to the onset of myelination, suggesting that axonal loss is not secondary to myelin defects. The loss of the axonal targets may have led oligodendrocytes to inappropriately myelinate neuronal cell bodies. Consistent with this, GANT61, which reduces oligodendrocyte number, caused a reduction in aberrantly myelinated neuron cell bodies in DomA-exposed fish. Together, these results suggest that DomA initially alters reticulospinal neurons and the loss of axons causes aberrant myelination of nearby cell bodies. The identification of initial targets and perturbed cellular processes provides a mechanistic understanding of how DomA alters neurodevelopment, leading to structural and behavioral phenotypes.

## INTRODUCTION

Harmful algal blooms (HABs) are a growing global threat that can lead to illnesses in humans and wildlife, socioeconomic losses, and disruptions to marine ecosystems.^1,2^ Some HABs can produce potent toxins that directly affect human health.^3–5^ Domoic acid (DomA) is a neurotoxin^6,7^ that is primarily produced by species in the diatom genus, *Pseudo-nitzschia*.^8,9^ Following the formation of toxigenic *Pseudo-nitzschia* blooms, DomA can accumulate in seafood, making it unsafe for consumption. The consumption of DomA-contaminated seafood at high doses leads to a syndrome called ‘Amnesic Shellfish Poisoning’ with symptoms that include memory loss, seizures, coma, and even death.^10–12^ To protect people against this syndrome, a regulatory limit has been set at 20 μg of DomA per g shellfish tissue.^13,14^ This limit was designed to protect adults from acute toxicity. However, mounting evidence suggests that developmental exposures to doses of DomA that do not cause overt symptoms can lead to lasting changes in tissue morphology,^15–17^ neural activity patterns,^18^ and behavior.^18–26^ The cellular targets and initiating molecular events that underlie these changes are largely unknown.

Our previous work has shown that DomA targets both myelin in the spinal cord and reticulospinal neurons – hindbrain neurons with axons that extend to the spinal cord.^27,28^ However, it is unclear whether DomA initially targets axons, leading to deficits in myelination, or whether DomA initially targets myelinating oligodendrocytes, leading to later axonal degeneration. To resolve this, we assessed the effects of DomA on these potential cellular targets close to the time of exposure to determine which cell types are the first to be perturbed by DomA.

DomA-induced disruptions to myelin sheath formation in the central nervous system^27,28^ led us to look into the effects of DomA on cells of the oligodendrocyte lineage, which are responsible for myelinating the central nervous system.^29,30^ Successful myelination requires oligodendrocyte precursor cells (OPCs) to form, migrate, differentiate into oligodendrocytes, and then initiate the process of myelination in different regions within the nervous system.^31,32^ Both OPCs and mature oligodendrocytes express functional ionotropic glutamate receptors, making them potential cellular targets for DomA.^33–37^ Previous studies have shown that kainate, a structural analog of DomA, causes cell death in OPC cultures at concentrations comparable to those affecting neurons.^38–41^ Furthermore, binding and activation of AMPA receptors inhibits the proliferation and differentiation of OPCs into mature oligodendrocytes.^35,42^ While generally less sensitive to kainate-induced cell death,^40^ mature oligodendrocytes have also been shown to undergo demyelination after chronic direct infusion of kainate on the optic nerves.^43^ All of this suggests that developmental exposure to DomA may target the oligodendrocyte lineage, and that exposure during early development may inhibit OPC differentiation, disrupt myelin sheath formation, and lead to cell death in the oligodendrocyte lineage.

The objective of this study was to identify the initial cellular targets of DomA. To accomplish this, we exposed fish to DomA at 2 dpf as before^27,28^ and then assessed the effects on candidate cell targets between 4 and 8 hours post-exposure, prior to myelination. Candidate cell targets included OPCs, reticulospinal neurons, sensory neurons, and motor neurons. We also further characterized DomA’s effects on the oligodendrocyte-lineage cells that are responsible for myelination by quantifying these cells and characterizing the myelination capacity of individual oligodendrocytes. The results identify reticulospinal neurons as the initial cellular targets for DomA and further characterize how the loss of their axons may have led to aberrant oligodendrocyte development and function.

## METHODS

### Fish husbandry and lines used

These experiments were approved by the Woods Hole Oceanographic Institution Animal Care and Use Committee (Assurance D16-00381 from the NIH Office of Laboratory Animal Welfare). Embryos were maintained at 28.5°C with a 14:10 light dark cycle during the experimental period in 0.3x Danieau’s medium. The following transgenic lines were used: *Tg(cntn1b:EGFP-CAAX)*,^27^ *Tg(mbp:EGFP-CAAX)*,^44^ *Tg(sox10:RFP)*,^45^ *Tg(nkx2.2a:mEGFP)*,^46^ *Tg(sox10:mRFP)*,^47^ and *Tg(mbp:EGFP)*.^48^

### Domoic acid exposure paradigm

Preparation of domoic acid solutions and exposure of embryos were described earlier (Panlilio et al., 2020). Briefly, domoic acid (Sigma-Aldrich, MO) was dissolved in a diluted embryo medium (0.2x Danieau’s) to generate stock concentrations of 0.676 μg/μl and 1.4 μg/μl (10 μl) that were stored at −20°C. Working solutions were prepared fresh prior to microinjection. Microinjections were performed using IM-300 microinjector that was calibrated to deliver 0.2 nL of solution.

DomA (nominal dose of 0.14 ng) was intravenously microinjected into the common posterior cardinal vein at 48-53 hpf ^49^ while controls from the same breeding cohort were injected with the saline vehicle (0.2x Danieau’s). To perform intravenous microinjections, fish were anesthetized with 0.10% tricaine methanesulfonate (MS222) and placed laterally on dishes coated with 1.5% agarose. An injection was deemed successful if there was a visible displacement of blood cells. Fish that showed evidence of being incorrectly injected were immediately removed from the study.

### Live imaging with confocal microscopy

Transgenic fish were live imaged using confocal microscopy. To accomplish this, fish were anesthetized in MS222 (0.16%), and then embedded laterally in 1-1.5% low melt agarose in glass bottom microscopy dishes. Fish were then imaged on the confocal microscope (LSM 710 or LSM 780) with the 40x water immersion objective (C-Apochromat, 40x, NA 1.1) or a 20x air objective (Plan-Apochromat, 20x, NA 0.8).

### OPC cell counts

Two double transgenic lines were used to assess potential effects of DomA on OPC counts – (*Tg(olig2:EGFP x Tg(sox10:mRFP)*) and (*Tg(nkx2.2a:mEGFP)* x *Tg(sox10:RFP)*). Following DomA exposure at 2 dpf, fish were laterally mounted at approximately 2.5 dpf, and their anterior spinal cords were imaged (roughly within the region from somites 8-12). Imaging files were blinded (treatment information was removed from the file). For each imaging file, the OPC cell bodies within the dorsal spinal cord were manually quantified by generating 3D projections on ImageJ, then rotating the stacks along the x-axis. The (*Tg(olig2:EGFP x Tg(sox10:mRFP)*) line was used to quantify OPCs and pre-myelinating oligodendrocytes in the dorsal spinal cord (cells that had membranes labeled with RFP and cell bodies labeled with EGFP). The (*Tg(nkx2.2a:mEGFP)* x *Tg(sox10:RFP)*) line was used to quantify the subset of OPCs that were fated to differentiate into myelinating oligodendrocytes (cells that had membranes labeled with EGFP and cell bodies labeled with RFP).

Experiments were performed twice (Supplemental Fig. 1A and 1B). The two experimental trials using *Tg(nkx2.2a:mEGFP)* x *Tg(sox10:RFP)* fish yielded significantly different OPC counts, as determined by a negative binomial model with experimental trial and treatment (control vs. DomA-exposed) as predictors using the ‘glm.nb’ function of the MASS package in R.^50^ The second trial had fish with an average of 1.6 times more OPCs than fish in the first trial, regardless of treatment group (CI = [1.46 1.74], p = < 2e-16, Supplemental Fig 1B). To address this, DomA was compared to controls within the same experiment.

### Myelinating oligodendrocyte cell counts

Similar procedures were used for counting myelinating oligodendrocytes. Transgenic fish that expressed GFP in myelinating cells, *Tg(mbp:EGFP)*, were laterally mounted at 4 dpf, and their anterior spinal cords were imaged around somites 6-10. Imaging files were then blinded, and myelinating oligodendrocytes were counted by rotating image stacks along the x-axis. To determine whether DomA alters the number of myelinating oligodendrocytes, a negative binomial model was constructed with dose (0, 0.14, or 0.18 ng of DomA) as a categorical predictor using the ‘glm.nb’ function of the MASS package in R.^50^

Cytoplasmic localized GFP also faintly labels myelin sheaths, allowing us to qualitatively classify myelin defects in addition to quantifying the number of oligodendrocytes for every individual fish. To see whether there was a correlation between oligodendrocyte number and severity of myelin defect, a negative binomial model was constructed with the myelin classification as predictors for each trial separately. To combine data from multiple experimental trials, a negative binomial regression model with random effects was used, with repeat trials taken into account as random factors (glmer.nb(), lme 4 package, R). One sample was excluded from the analysis as it did not fit the four categories.

### Mosaic labeling of myelin sheaths from individual oligodendrocytes

To determine whether DomA alters myelin sheath length and number, we performed mosaic labelling by injecting an *mbp:EGFP-CAAX* construct at the 1-4 cell stage. The *mbp:EGFP-CAAX* construct was generated by using *Tg(mbp:EGFP-CAAX)* genomic DNA as a template and amplifying the region that contained the *mbp* promoter along with the membrane bound EGFP.^51,52^ The sequence was then placed into the pkHR7 plasmid containing ISCe-1 restriction sites.^53^ Embryos were exposed to either DomA or the saline vehicle and those with sparsely labeled oligodendrocytes were imaged at 4 dpf. Some embryos had multiple labeled oligodendrocytes. Oligodendrocytes from the same fish were assigned a ‘fish ID’ number to distinguish multiple oligodendrocytes that came from the same fish from oligodendrocytes that came from different fish.

Images of individual oligodendrocytes were blinded prior to image analysis. To perform the analysis, myelin sheaths from individual oligodendrocytes were traced using the ImageJ plugin Simple Neurite Tracer.^54^ For each individual oligodendrocyte, the total number of myelin sheaths and the number of circular profiles were counted and the length of the individual myelin sheaths was approximated. To determine whether there was an effect of treatment on the average length of the myelin sheaths per oligodendrocyte, a nonparametric multivariate response analysis was performed (art(), ARTool R package).^55^ To account for potential variations in responses from different individual fish, fish ID numbers were included in the model as random effects.

To determine whether DomA altered the number of myelin sheaths or the number of circular myelin membranes, a generalized linear model with a negative binomial distribution was built (glmer.nb(), lme4 R package), assigning treatment (Control or DomA) and fishID as fixed effects.

### DomA effect on primary motor neuron axons

*Tg(olig2:EGFP)* x *Tg(sox10:mRFP)* embryos were used to assess the effects of DomA on primary motor neurons. The presence or absence of the main axons for three primary motor neuron classes were noted for each fish. Following this classification, quasi-binomial logistic regression analyses were performed to determine whether exposure to DomA led to significant differences in the presence of main axons in the three types of primary motor neurons.

### Immunohistochemistry

Fish used for imunohistochemical analyses were exposed to 75 μM 1-phenyl-2-thiourea (PTU) at 24 hpf and onwards to inhibit pigment formation and to allow for imaging of axonal structures within different brain regions.^56^

Embryos were anesthetized and then fixed in 4% paraformaldehyde overnight at 4°C. Antigen retrieval was done by placing tissue in 150 mM Tris Hcl (pH 9.0) in a 70°C water bath for 15 minutes.^57^ Tissues were processed as whole mounts and permeabilized using ice-cold acetone (7 minutes). Samples were then blocked in 10% normal goat serum, 1% BSA and 1% DMSO, followed by 1-3 day incubations in primary antibodies (α-neurofilament associated antigen (3A10) −1:100 dilution, Developmental Studies Hybridoma Bank; α-acetylated tubulin – 1:500 dilution, Santa Cruz Biotechnology). After several washes with phosphate buffered saline (1X PBS, pH 7.2 - 7.4), samples were incubated in secondary antibodies (1:400 Alexa Fluor 488 Goat α-mouse or Alexa Fluor 596 Goat α-mouse; Abcam). Samples were then placed in antifade mountant (Prolong or SlowFade Diamond mountant, Invitrogen) between bridged #1.5 coverslips for imaging.

Embryos stained with 3A10 were mounted dorsally to image the Mauthner cell bodies and hindbrain and midbrain axonal tracks. Embryos stained with α-acetylated tubulin were mounted laterally to assess the sensory neuron ganglia and lateral line structures; these embryos were a subset of those imaged previously^27^ in a different anatomical region (trunk). Following antibody staining and mounting, fixed larvae were imaged on the confocal microscope using either 20x air objective (Plan-Apochromat 20x NA 0.8) or the 40x water immersion objective (C-Apochromat, 40x, NA 1.1).

### DomA effect on Mauthner cells

Following 3A10 staining and confocal imaging, the number of Mauthner cells (0, 1, or 2) was noted for each fish. To determine whether DomA alters Mauthner cell number, ordinal logistic regression analyses were done (polr(), MASS R package), with treatment (DomA vs. control) as a categorical predictor.

### Characterization of circular myelin features

*Tg(cntn1b:EGFP-CAAX)* x *Tg(sox10:mRFP)* are double transgenic fish that express RFP in myelin sheaths and EGFP in the membranes of specific neurons. After exposure to DomA or vehicle, larvae were imaged at 5 dpf. Image stacks were then examined to see whether the circular oligodendrocyte membranes labeled with RFP surrounded neuronal cell bodies labeled in EGFP. The larvae imaged were a subset of those imaged previously^27^ in a different anatomical region (cranial region).

### Scanning Electron microscopy

Tg(mbp:EGFP-CAAX) fish were exposed to 0.14 ng of DomA (nominal dose) at 2 dpf. At 5 dpf, fish were screened using the widefield epifluorescence microscope for myelin defects. DomA-exposed fish with a high prevalence of circular membranes were preselected for electron microscopy. Control fish with normal-looking myelin were also selected.

The electron microscopy work, including fixation, embedding, and thin sectioning, were done by the Marine Biological Laboratory Central Microscopy facility (Woods Hole, MA). Using a modified rOTO (reduced osmium—thiocarbohydrazide—osmium) protocol, larvae were fixed in a 2% paraformaldehyde and 2.5% glutaraldehyde in 0.2M cacodylic buffer with 3% sucrose for 1.5 hours. Following three rinsed in 0.2M cacodylic buffer, samples were postfixed in 1% osmium tetroxide and 0.75% potassium ferricyanide in 0.2M cacodylic buffer for 40 minutes. Samples were then rinsed in distilled water and incubated in 0.25% thiocarbohydrazide for 10 min. Following another rinse, they were embedded in agarose and post-treated with 0.5% osmium tetroxide for 30 minutes, rinsed and incubated overnight in 1% uranyl acetate at 4°C. Thoroughly rinsed in distilled water samples were then dehydrated in graded ethanol solutions and embedded in epoxy resin following standard protocol. All chemicals and materials were purchased from Electron Microscopy Sciences (Hatfield, PA).

For SEM imaging, sections were cut ~70nm thick and collected on a continuous feed of Kapton tape using an RMC Boeckleler ATUMtome ultramicrotome. Carbon coated series of sections mounted on a silicon wafer were imaged using SEM Zeiss Supra40VP. Whole spinal cords were imaged at 10 nm and 5 nm per pixel resolution in one domoic acid treated fish to screen for neuronal cell bodies with myelin surrounding them over roughly 50 slices in the medial spinal cord. When areas of interest were identified, smaller regions were imaged at 5 nm resolution using SEM.

The TEM imaging was then done using thin sections that were ~70nm thick cut and collected on copper grids using an RMC Boeckleler ATUMtome. Sections poststained with 1% uranyl acetate and 0.04% lead citrate were imaged using TEM microscope JEOL-JEM-200CX.

### GANT61 pharmacology

GANT61 (2,2’-[[dihydro-2-(4-pyridinyl)-1,3(2H,4H)-pyrimidinediyl]bis(methylene)]bis[N,N-dimethyl-benzenamine) is a GLI antagonist that was used to reduce the number of oligodendrocytes.^58,59^ GANT61 (Sigma-Aldrich, MO) was dissolved in 100% DMSO to create 10, 25, and 40 mM (1000x) stock solutions. Working solutions were prepared fresh prior to the experimental trials. Double transgenic fish (*Tg(mbp:EGFP)* x *Tg(mbp:EGFP-CAAX)*) were used for these experiments to distinguish aberrant circular membrane profiles (which only had labeled membranes) from oligodendrocyte cell bodies (which had both labeled cytoplasm and the membranes). These double transgenic fish were exposed to different concentrations of GANT61 (0, 10, 25, or 40 μM) in rearing medium (0.3X Danieu’s) from 1.5-4 dpf, and a subset of fish were exposed to either 0.14 ng of DomA or saline vehicle at 2 dpf. Larvae were then live imaged at 4 dpf using the confocal microscope (LSM 710) using the techniques outlined above.

The number of oligodendrocyte cell bodies in the dorsal spinal cord was determined. GANT61 treatment in both control and DomA-treated fish led to the labeling of several cells in the most dorsal regions of the spinal cord that could not be identified. Additionally, the large myelin sheaths present in the ventral region of the spinal cord occluded some oligodendrocyte cell bodies, making it difficult to accurately quantify the oligodendrocytes in this region. Given these two considerations, we simplified the analysis by only quantifying oligodendrocytes and circular myelin features in the dorsal region of the spinal cord. To obtain an estimate of the number of ectopically myelinated cell bodies per oligodendrocyte, we divided the total number of these cell bodies by the number of oligodendrocytes found within the total imaging area. To assess the effect of treatment on the average number of cell bodies per oligodendrocyte an Analysis of Variance (ANOVA) followed by Dunnett-type contrast was performed (multicomp, stats R libraries).^60^

### Statistical analyses of data from GANT61-treated fish

To identify the effects of GANT61 treatment on the number of oligodendrocytes and circular bodies, negative binomial regression models were used (glm.nb()). To assess the potential correlation between oligodendrocyte counts and circular cell body counts, a spearman correlation was performed (cor.test(), stats R package).

## RESULTS

### Oligodendrocyte precursor cell number is unaltered by DomA exposure

Previous work has shown that exposure to DomA at 2 dpf led to myelin deficits along with the appearance of unusual circular profiles.^28^ To determine whether myelination deficits were driven by a decrease in the supply of oligodendrocyte precursor cells (OPCs) or myelinating OPCs, we quantified these cells in DomA-exposed larvae using two double transgenic lines (Fig. 1A, 1D).

**Figure 1:**
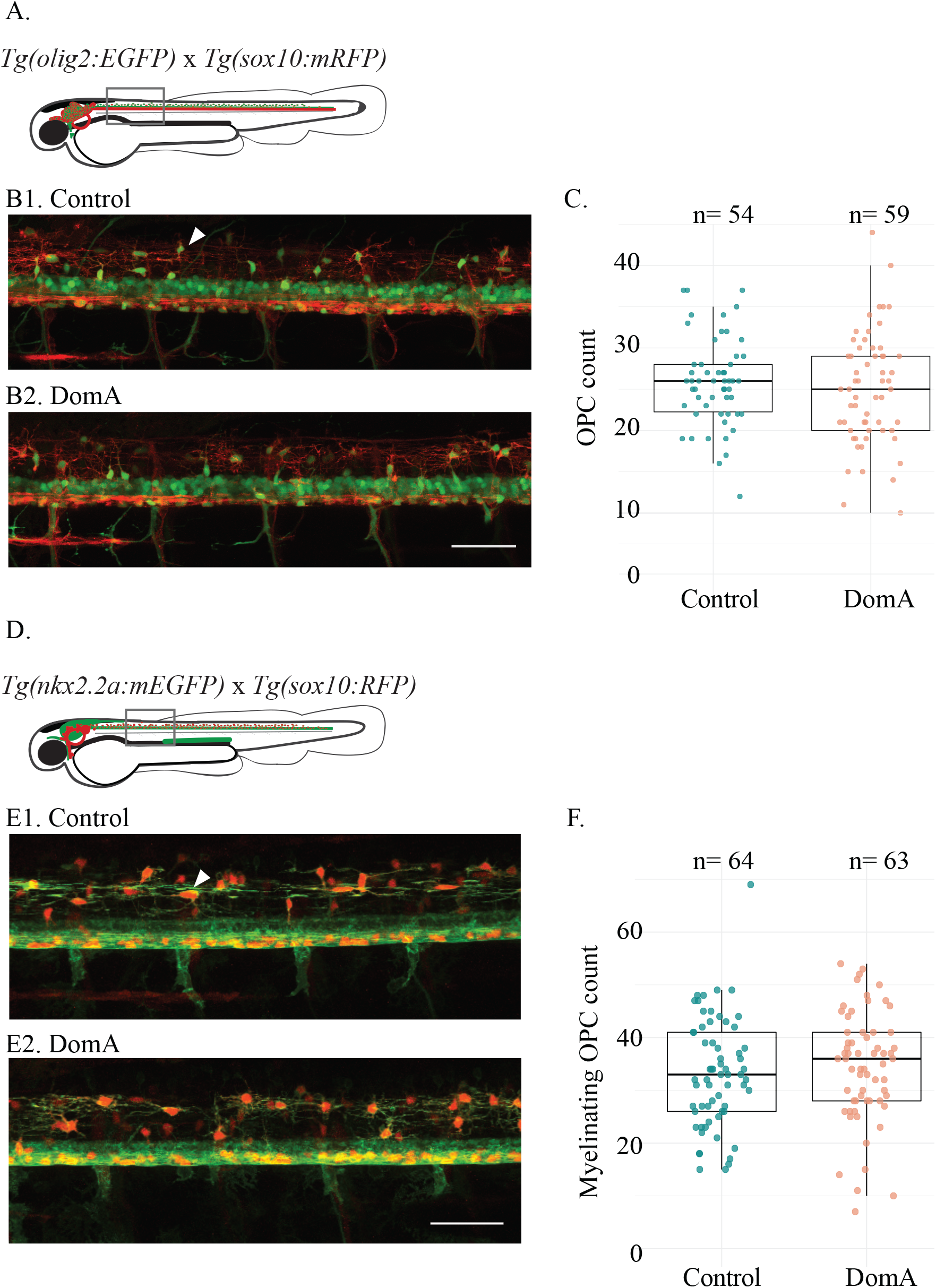
Exposure to DomA at 2 dpf did not reduce the number of oligodendrocyte precursor cells prior to myelination. (A) Diagram of laterally mounted *Tg(olig2:EGFP)* x *Tg(sox10:mRFP)* double transgenic larvae. The rectangle delineates the approximate location within the spinal cord where the image was acquired. (B) Representative images of the double transgenic fish, *Tg(olig2:EGFP)* x *Tg(sox10:mRFP)*, imaged at 2.5 dpf. Control (B1) and DomA-exposed fish (B2). White arrow points to a dorsal OPC. Scale bar = 50 μm (C) Oligodendrocyte precursor cell (OPC) count in dorsal spinal cords of the control and DomA-exposed double transgenic fish —*Tg(olig2:EGFP)* x *Tg(sox10:mRFP)*. Each point represents the number of OPCs counted within the 403.1μM imaging area in a single fish (Control — median = 26 and IQR = 6; DomA — median = 25, IQR = 9). (D) Diagram of laterally mounted *Tg(nkx2.2a:mEGFP)* x *Tg(sox10: RFP)* double transgenic larvae. The rectangle delineates the approximate location within the spinal cord where the image was acquired. (E) Representative images of the double transgenic fish, *Tg(nkx2.2a:mEGFP)* x *Tg(sox10:RFP)*, imaged at 2.5 dpf. Control (E1) and DomA exposed fish (E2). White arrow points to a dorsal OPC in the myelinating lineage. Scale bar = 50 μm (F) Counts of OPCs that are fated to be myelinating oligodendrocytes in the dorsal spinal cord in double transgenic fish — *Tg(nkx2.2a:mEGFP)* x *Tg(sox10: RFP)*. Each point represents the number of OPCs counted within the 354.3μM imaging area in a single fish. (Control – median = 33, IQR= 15, DomA – median = 36, IQR = 13). Data shown are from two combined trials. See Supplemental Fig. 1 for data from individual trials.

All OPCs were quantified using the (*Tg(olig2:EGFP)* x *Tg(sox10:mRFP)*) line (Fig. 1A). Exposure to DomA did not reduce the total number of OPCs in the ventral spinal cord prior to myelination (Fig. 1B, 1C). To determine whether DomA selectively targets the subset of OPCs that are fated to become myelinating oligodendrocytes *(myelinating OPCs)*, we used a double transgenic line (*Tg(nkx2.2a:mEGP)* x *Tg(sox10:RFP)*) (Fig. 1D). As with total OPCs, there were no differences in the number of myelinating OPCs between control and DomA-treated larvae (Fig. 1E, 1F).

### DomA exposure leads to a reduction of myelinating oligodendrocytes in a dose-dependent manner

OPCs differentiate into myelinating oligodendrocytes. We used the *Tg(mbp:EGFP)* line to determine the effect of DomA exposure at 2 dpf on the number of myelinating oligodendrocytes at 4 dpf (Fig. 2A). DomA reduced the number of myelinating oligodendrocytes in a dose-dependent manner. Exposure to 0.14 ng DomA led to a 12.9% decrease relative to controls, while 0.18 ng DomA led to a 31.8% decrease in the number of oligodendrocytes relative to controls (0.14 ng DomA -- Coefficient = −0.130, Error = 0.031, p = 2.37 e-5; 0.18 ng DomA -- Coefficient = −0.409, Error = 0.046, p= < 2 e-16) (Fig. 2B; Supplemental Fig. 2).

**Figure 2:**
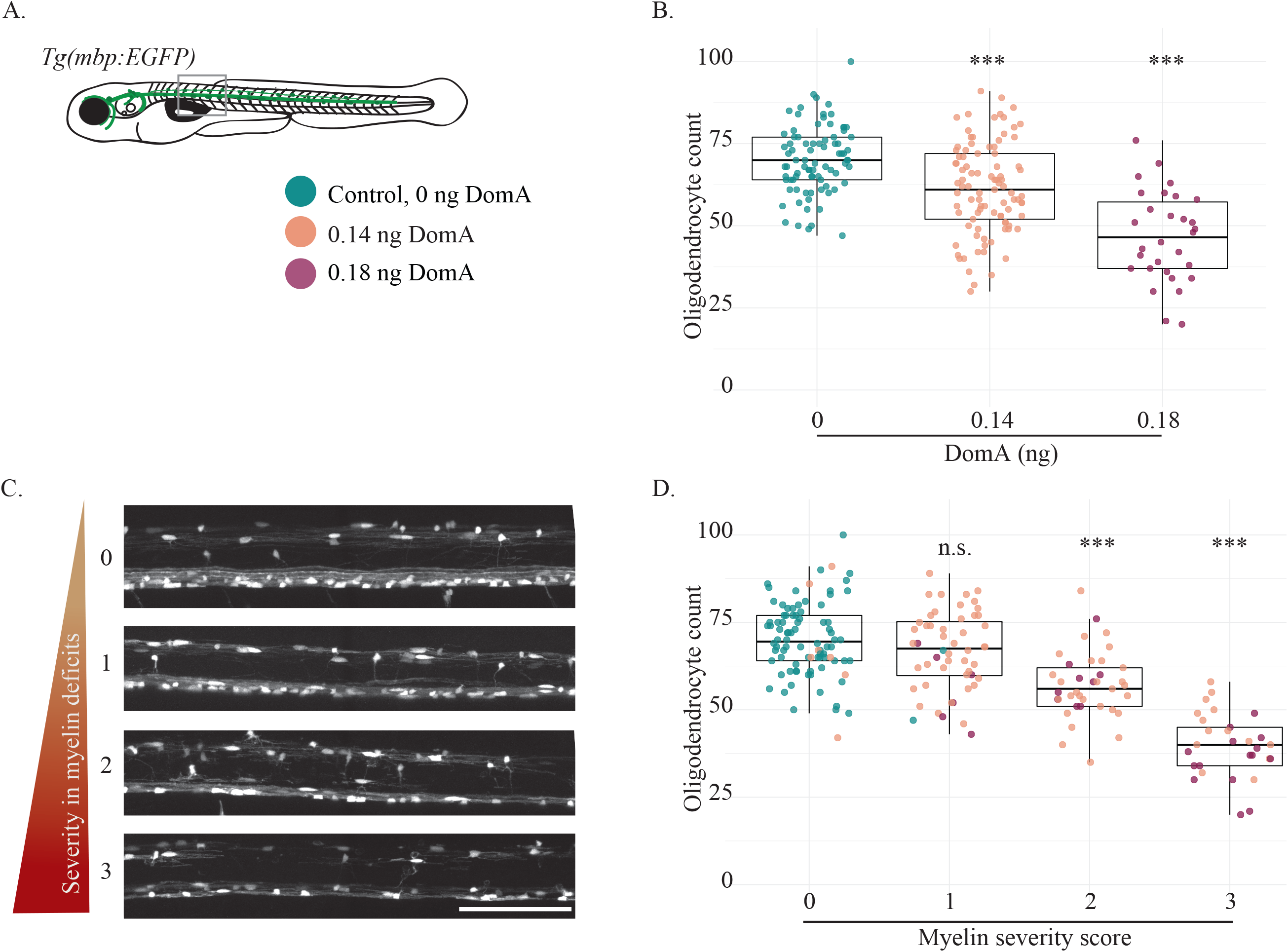
Exposure to DomA at 2 dpf reduces the number of myelinating oligodendrocytes in fish with severe myelin defects, and in a dose dependent manner. (A) The *Tg(mbp:EGFP)* transgenic line was used to quantify myelinating oligodendrocytes at 4 dpf. The rectangle delineates the approximate location within the spinal cord where the image was acquired (somites 6-10). (B) Number of myelinating oligodendrocytes quantified at 4 dpf in the spinal cords of fish exposed to different doses of DomA at 2 dpf. Each point represents the number of myelinating oligodendrocytes within the 403.9μM imaging area in a single fish. (Control (0 ng) – median = 70 and IQR = 13; DomA 0.14 ng – median = 62, IQR = 20; DomA 0.18 ng – median = 47, IQR = 20). (C) Representative images of laterally mounted *Tg(mbp:EGFP)* fish classified by myelin severity, ranging from 0, representing control-like myelin sheaths, to 3, representing the most severe myelin phenotype observed. (D) Fish used in Fig 2B were further subdivided by the severity of the myelin defect observed. The myelinating oligodendrocyte counts were then plotted against the myelin severity score. Scale bar = 100 μm, n.s. = not significant, *** p < 1 e −3 using a generalized mixed effects models with a negative binomial distribution

To determine the relationship between the severity of myelin defects and the number of oligodendrocytes present, individual fish from these experiments were also classified by the severity in myelin defects observed (Fig. 2C). Fish that had the least severe myelin defects (category 1) did not have significantly different oligodendrocyte numbers compared to fish with no myelin defects (category 0) (Coefficient = −0.05, Error= 0.03, p = 0.057) (Fig. 2D). This suggests that reductions in oligodendrocyte numbers are not necessary for the appearance of the least severe myelin phenotypes. In contrast, fish with higher myelin severity phenotypes (category 2 and 3) had significantly reduced oligodendrocyte numbers. Fish with category 2 phenotypes had an 18% reduction in oligodendrocytes compared to controls (Coefficient = −0.195, Error =0.032, p = 1.35 e-9), while fish with most severe defects (category 3) had the lowest oligodendrocyte numbers (42% reduction relative to controls (Coefficient = −0.544, Error = 0.039, p = < 2e-16)) (Fig. 2D). These results demonstrated that exposure to DomA, especially at the higher dose, reduced the number of myelinating oligodendrocytes and that the extent of reduction was related to the severity of the myelin phenotypes observed. However, there were instances when exposure to DomA resulted in less-severe myelin phenotypes and that did not result in the reduction in oligodendrocyte number, suggesting that other effects in addition to the reduction in oligodendrocyte number may contribute to the reduced myelination.

### Individual oligodendrocytes produce aberrant myelin sheaths following exposure to DomA

Myelin deficits could also result from the inability of individual oligodendrocytes to myelinate axons appropriately. To address this, we sparsely labeled oligodendrocytes and counted myelin sheath length and number for individual oligodendrocytes (Fig. 3A). Oligodendrocytes from DomA-exposed larvae produced significantly fewer myelin sheaths (Coefficient = −1.01, Std. Error = 0.11, p < 2 e-16) that were also shorter relative to controls (F(1, 141.25)= 64.101, p= 3.99e-13) (Fig. 3B, 3C). In addition to the myelin deficits, DomA-exposed fish also had a higher number of unusual circular profiles instead of the characteristic long and thin myelin sheaths (Coefficient = 2.52, Std. Error = 0.294, p = 1.99e-14) (Fig. 3D).

**Figure 3:**
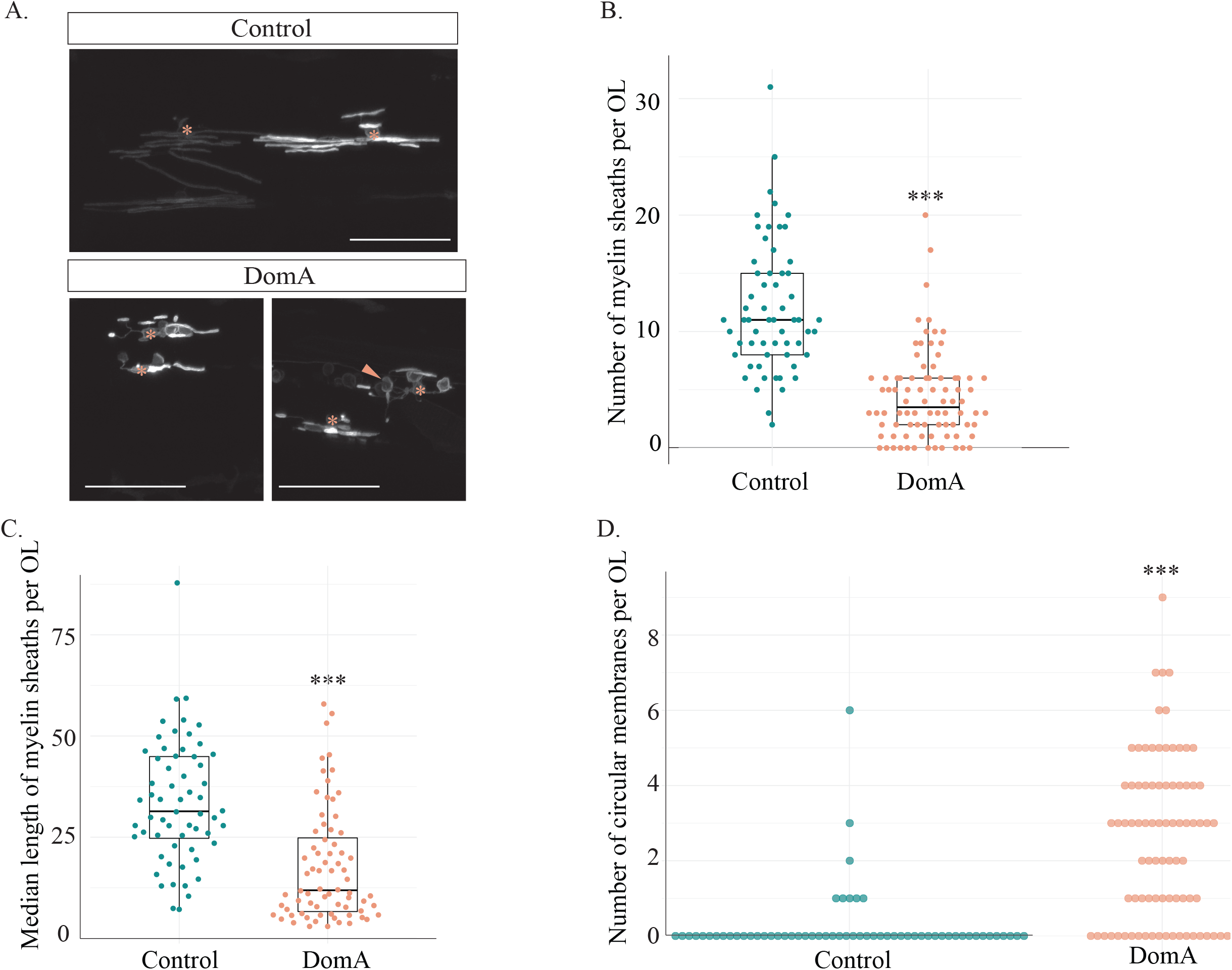
DomA reduces the length and number of myelin sheaths produced by individual oligodendrocytes by 4 dpf. (A) Representative images of mosaically labeled oligodendrocytes in DomA-exposed (2 dpf injected) and control fish following 1 cell injections of the reporter construct, mbp:EGFP-CAAX into *Tg(sox10:RFP)* background. Only EGFP channel is presented to show the sparsely labeled myelin sheaths. Asterisks mark oligodendrocyte cell bodies. Peach arrow points to aberrant circular profiles. Scale bar = 50 μm (B) Number of myelin sheaths produced by single oligodendrocytes (OL) in both control and DomA exposed fish. (C) Median length of the myelin sheaths produced by individual OLs in μm. OL without myelin sheaths were excluded from this graph. (D) Number of circular myelin membranes produced by individual OLs.

### DomA exposure at 2 dpf, but not at 1 dpf, leads to the loss of the Mauthner neurons prior to myelination

Aberrant myelination may result from both the loss of myelinating oligodendrocytes and the inability of the remaining oligodendrocytes to myelinate axons. However, the effects of DomA on the oligodendrocytes do not rule out the possibility that DomA leads initially to the loss of neurons and their axons, which subsequently has secondary effects on the oligodendrocytes. To determine whether DomA first targets the axons prior to myelination, we performed immunohistochemistry and live imaging of specific neuronal populations prior to myelination.

Many of the reticulospinal neurons, including the Mauthner cells, were present by 52 hpf, which was within the 2 dpf (48-53 hpf) injection exposure period (Fig. 4A,B). However, by 60 hpf (8-12 hours post exposure), both Mauthner neurons were lost in the majority of fish exposed to DomA (40/72) (Fig. 4C, Fig. 4D, p < e-13; Supplemental Fig. 3). In contrast, both Mauthner neurons were found in all control fish tested (76/76). Since Mauthner neurons were present at the time of exposure, DomA did not inhibit the formation of the Mauthner neurons but rather led to the loss of these neurons.

**Figure 4:**
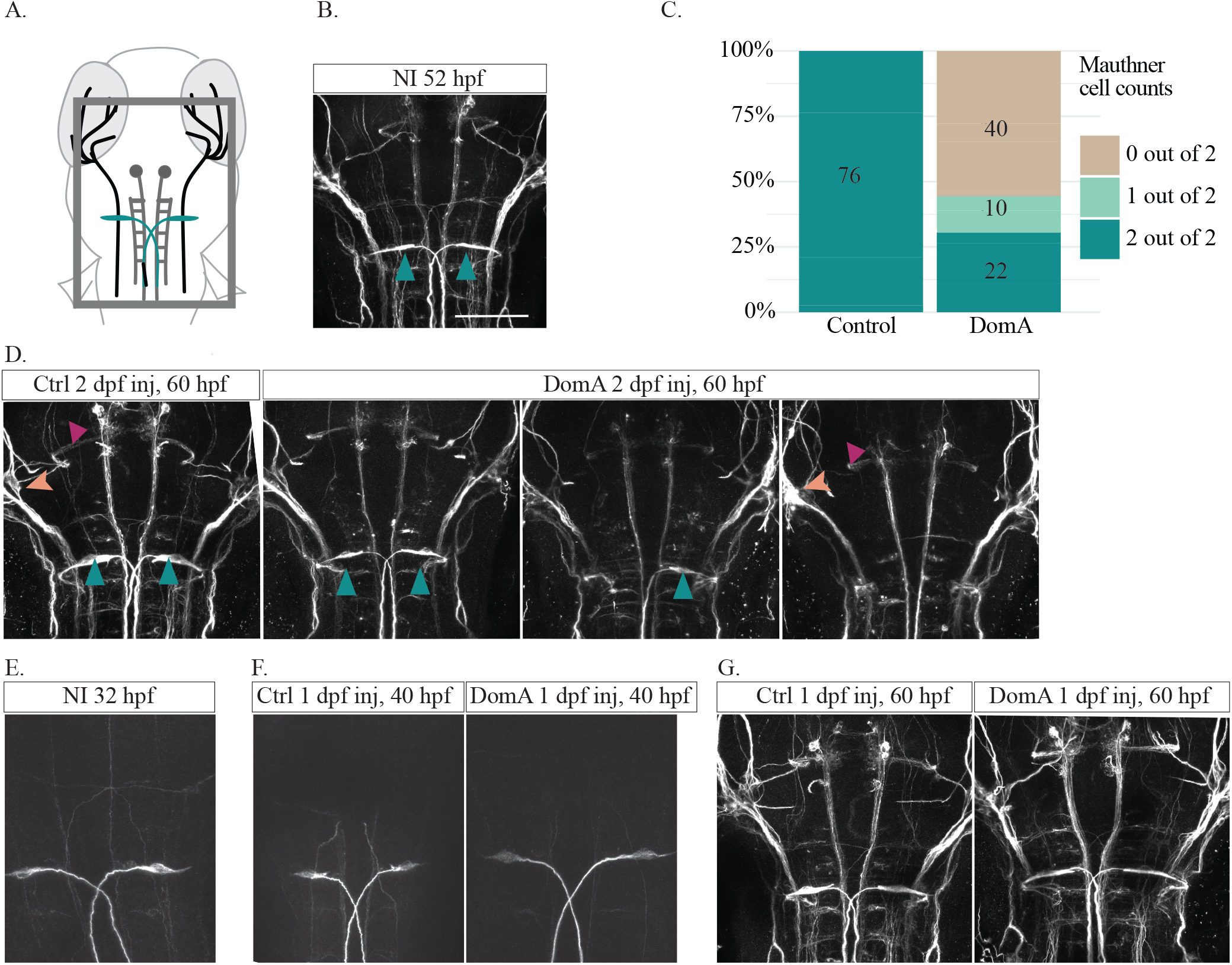
Exposure to DomA at 2 dpf (but not 1 dpf) leads to the loss of the Mauthner neuron prior to myelination. (A) Diagram of 2.5 dpf embryos that were dorsally mounted and immunostained with anti-3A10. The rectangle delineates the approximate area that was imaged. Mauthner cells are labeled in teal. (B) 3A10 immunostaining of 52-hpf, non-injected (NI) embryos confirmed that all the major reticulospinal neurons are present by the 2 dpf (48-53 hpf) injection period (n= 26). Teal arrows label Mauthner cells. Scale bar = 100 μm. (C) Percentage of embryos exposed to DomA (2 dpf) or vehicle that had 0, 1, or 2 Mauthner cells after being stained for anti-3A10 at approximately 60 hpf. Numbers of larvae with each phenotype are listed within each bar. Data were aggregated from four trials. See Supplemental Fig. 3 for individual trial data. (D) 3A10 immunostaining of embryos that were exposed to DomA at 2 dpf, then processed approximately 8 hours post exposure (60 hpf). Teal arrows label Mauthner cells, peach arrows label the anterior lateral line, and magenta arrows label the medial longitudinal fasciculus. Scale bar = 100 μm. (E) 3A10 immunostaining of 32-hpf, non-injected (NI) embryos confirmed that all the major reticulospinal neurons are present by 1 dpf injection period (n= 10). Scale bar = 50 μm. (F) 3A10 immunostaining of embryos that were exposed to DomA at 1 dpf (32 hpf), then processed approximately 8 hours post exposure (40 hpf). Control embryos (n=38); DomA exposed embryos (n= 38), with n= 37 fish with 2 Mauthner cells, and n= 1 with 1 Mauthner cell. Scale bar = 50 μm. (G) 3A10 immunostaining of embryos that were exposed to DomA at 1 dpf (32 hpf), then processed at 60 hpf. Control embryos (n= 35), DomA exposed embryos (n= 41 All control and DomA-exposed fish had 2 Mauthner cells). Scale bar = 50 μm.

Previous studies showed that exposure at 2 dpf (0.14 ng, injected at 48-52 hpf) led to startle response deficits and myelin defects, whereas exposures at 1 dpf (injected at 28-32 hpf) did not lead to any of these effects.^27,28^ We sought to determine whether, as seen for the behavioral and myelin phenotypes, the loss of Mauthner cells does not occur after exposure at 1 dpf. Even though the Mauthner cells were present at the time of exposure (Fig. 4E), fish exposed to DomA at 1 dpf did not lose Mauthner cells by either 8 hours post exposure (40 hpf) or by 60 hpf, which are times when 2-dpf-injected fish were observed to have lost one or both Mauthner cells (Fig. 4 F, G).

### DomA exposure does not lead to any apparent effects in selected sensory system structures

Previous work established that exposure to DomA at 2 dpf led to no apparent changes in sensory system structures or to the main axon of the caudal primary motor neurons by 5 dpf.^27^ To rule out the possibility that DomA perturbs these neuronal subtypes early in development even if they eventually recover in the larval stages, we assessed whether exposure to DomA altered specific sensory and motor neuron structures at 2.5 dpf.

DomA exposure did not lead to any apparent deficits in sensory ganglia that comprise the anterior lateral line including the nADso, nVDI, nADb and nAVm ganglia (Fig. 5A,B), nor did it lead to any apparent deficits in the anterior lateral line or the medial longitudinal fasciculus prior to myelination, when compared to controls (Fig. 4D).

**Figure 5:**
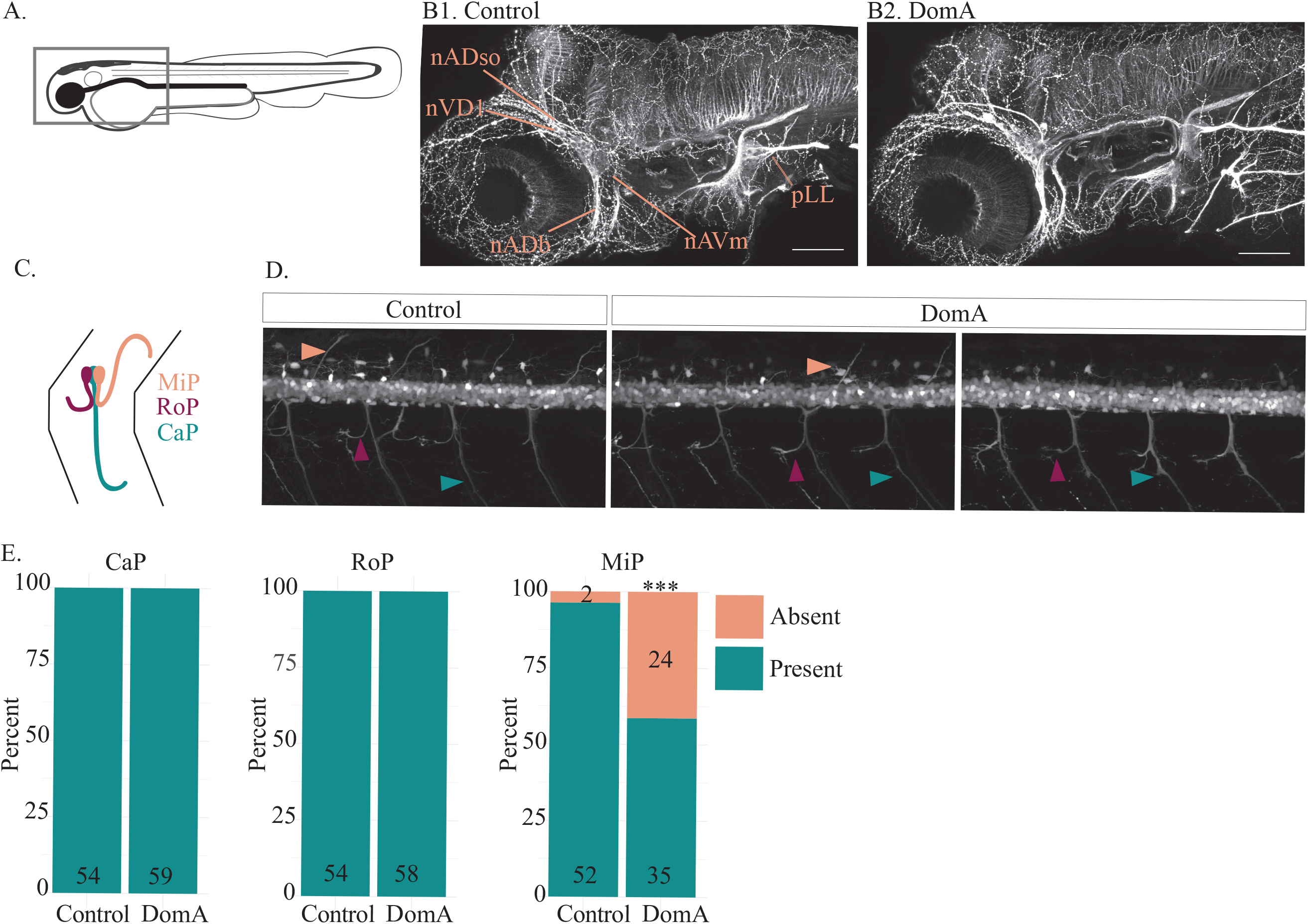
Exposure to DomA at 2 dpf does not alter selected sensory neuron structures or the main axons of two of the three primary motor neurons. (A) Diagram of 2.5 dpf embryos laterally mounted and immunostained with anti-acetylated tubulin. The rectangle delineates the approximate area that was imaged. (B) Representative images of control (n= 35) and DomA-exposed larvae (n=29)immunostained with anti-acetylated tubulin. Labels point to both the sensory ganglia that comprise the anterior lateral line (nADso, nVDI, nADb, nAVm) and to the peripheral lateral line (pLL). (C) Diagram of the axonal trajectories in the three types of primary motor neurons assessed in *Tg(olig2:EGFP)* fish. (D) Representative images of *Tg(olig2:EGFP))* x *Tg(sox10:mRFP)* fish exposed to vehicle or to DomA at 2 dpf then imaged at 2.5 dpf. Peach arrows point to the MiP main axons, magenta arrows point to the RoP main axons, and teal arrows point to the CaP main axons. Scale bar = 50 μm. These fish were the same fish used in Figure 1, with only the EGFP shown. (E) Presence or absence of the main axons of the three types of primary motor neurons in control and DomA-exposed fish. ***p < 0.001 Abbreviations: superior opthalmic ramus of the anterodorsal lateral line nerve (nADso), dorsolateral nerve of the trigeminal ganglion (nVDI), buccal ramus of the anterodorsal lateral line nerve (nADb) or the mandibular ramus of the anterior lateral line nerve (nAVm). Middle primary motor neuron (MiP), rostral primary motor neuron (RoP), Caudal primary motor neurons (CaP).

### DomA exposure alters the main axons in one of the three primary motor neuron classes

To determine whether DomA alters primary motor neurons at 2.5 dpf, prior to myelination, we used the *Tg(olig2:EGFP x Tg(sox10:mRFP)* line to assess the presence of the main axons of the caudal (CaP), middle (MiP), and the rostral (RoP) primary motor neurons (Fig. 5C-E). In agreement with previous studies, DomA did not alter the main caudal and rostral primary motor neurons (Fig. 5D,E). However, it significantly reduced the number of observed MiP primary motor neurons (Estimate = −2.91, Std. Error = 0.78, p = 0.0003) (Fig. 5D,E). Together, these results show that DomA does not alter selected sensory structures or CaP and RoP main primary motor neuron axons in embryos soon after DomA exposure — a finding that is consistent with effects previously described in the larval stages (5 dpf).^27^

### Domoic acid-induced axonal losses may contribute to observed myelin defects

DomA led to the loss of reticulospinal neurons and their axons prior to myelination (Fig. 4). The loss of subsets of hindbrain reticulospinal neurons and spinal interneurons would likely result in an overall decrease in axonal surface area. It is conceivable that changes to the axonal surface area could alter the spinal cord cellular environment and in turn lead to the myelin defects observed.

One of the characteristics of DomA-induced myelin defects is appearance of the unusual circular profiles rather than thin elongated myelin sheaths. Similar circular features have been characterized in a mutant for kif1 binding protein (*kif1bp*), which has 50-80% reductions in axon surface areas within the spinal cord. The circular features were identified as myelin that ectopically wrapped neuronal cell bodies.^59^ We thus sought to determine whether DomA-induced circular profiles were also ectopically myelinated neuronal cell bodies.

To exclude the possibility that the observed circular profiles were oligodendrocyte cell bodies, we used the double transgenic line, *Tg(mbp:EGFP-CAAX)* x *Tg(mbp:EGFP)*, to distinguish oligodendrocyte cell bodies, which are round but contain EGFP in the cytoplasm, from circular membranes, which are also round but contain EGFP only in their membranes. Control fish had labeled oligodendrocyte cell bodies but rarely had the hollow-looking circular features. In contrast, DomA-exposed fish had labeled oligodendrocyte cell bodies along with many of these circular features (Fig. 6A). These findings were also corroborated by experiments in which *Tg(sox10:RFP)* fish, which express RFP in all oligodendrocyte cell bodies, were injected with the mbp:EGFP-CAAX plasmid to mosaically express EGFP in myelin sheaths. Fish exposed to DomA showed numerous circular profiles that contained EGFP in the membrane with no RFP in their cytoplasm (Fig. 6B).

**Figure 6:**
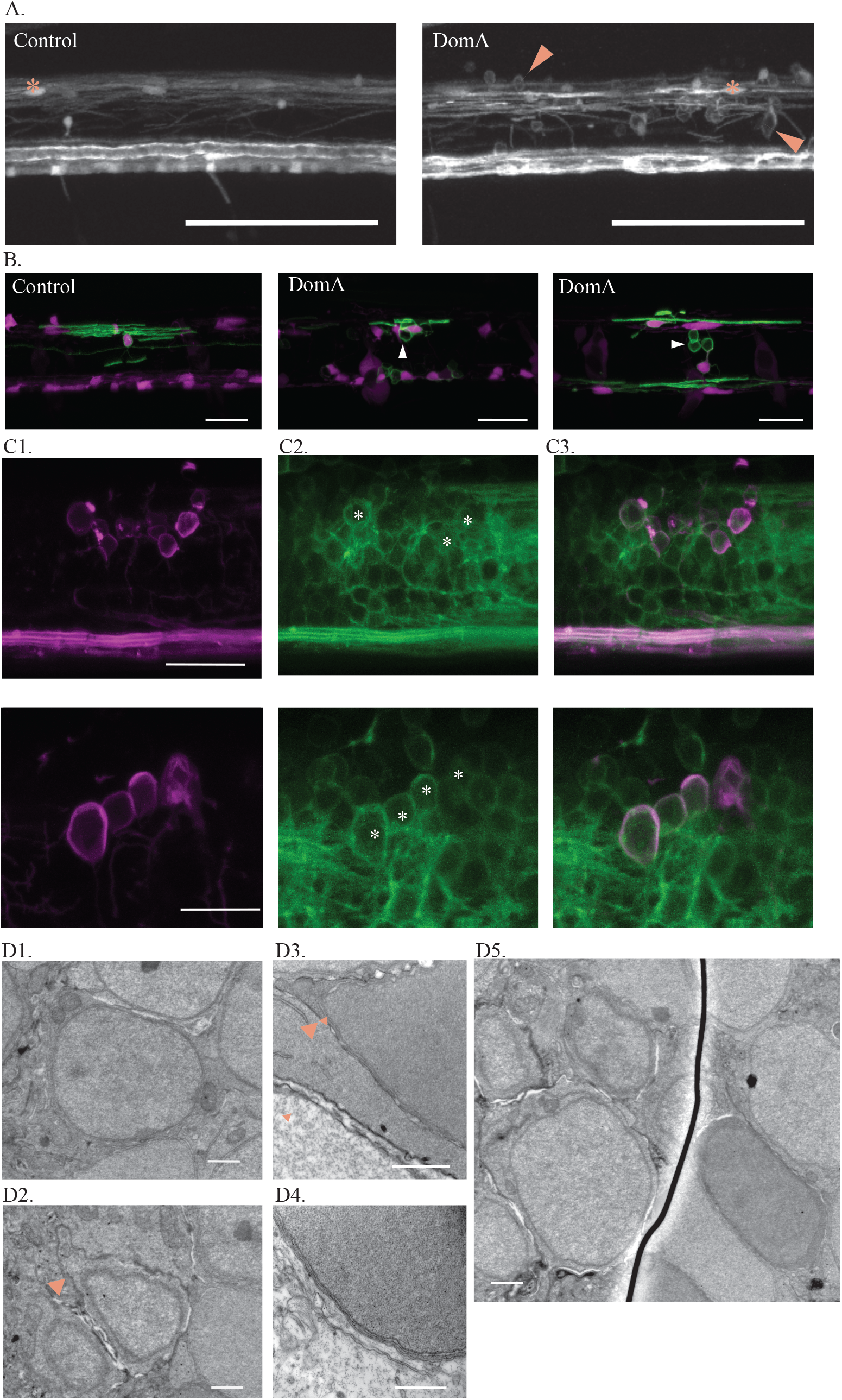
DomA exposure at 2 dpf leads to the formation of aberrant circular profiles that may be ectopically myelinated neuronal cell bodies. **(A)** *Tg(mbp:EGFP)* x *Tg(mbp:EGFP-CAAX)* double transgenic line labels both oligodendrocyte cell bodies and myelin sheaths. Asterisks mark oligodendrocyte cell bodies. Arrows label circular myelin membranes, which are commonly found in DomA-exposed larvae and rarely observed in controls. Scale bar = 100 μm. **(B)** Representative images of mosaically labeled oligodendrocytes in DomA-exposed (2 dpf) and control fish following 1 cell injections of the reporter construct, mbp:EGFP-CAAX into *Tg(sox10:RFP)* background, imaged at 4 dpf. Oligodendrocyte cell bodies are labeled in red. Arrows mark circular myelin membranes, which are outlined in green. These were the same fish used in Fig. 3, but with the red channel present to show the oligodendrocyte cell body. Scale bar = 25 μm. **(C)** *Tg(sox10:mRFP)* x *Tg(cntn1b:EGFP-CAAX)* double transgenic line imaged at 5 dpf. mRFP labels myelin sheaths and oligodendrocyte membranes. EGFP-CAAX labels the membrane of subpopulations of spinal cord neurons. Two examples of DomA-exposed double transgenic fish. Circular myelin membranes labeled in RFP (C1), neuronal cell bodies outlined in EGFP (C2). Images were then merged (C3). Asterisks mark neuronal cell bodies that are potentially associated with circular myelin membranes in the imaging plane. Scale bar = 20 μm. **(D)** Scanning electron micrograph of neuronal cell bodies in the spinal cords of *Tg(mbp:EGFP-CAAX)* fish. **(D1)** Example neuronal cell body in a control fish shows no evidence of myelin surrounding it. **(D2-D5)** Example neuronal cell bodies in DomA-exposed fish that may be wrapped by myelin. Arrows point to the putative myelin that surrounds the cell body. (n=1 control, n= 1 DomA-exposed) Scale bar= 1 μm.

To determine whether these circular features were ectopically myelinated neuronal cell bodies, we used a double transgenic line (*Tg(cntn1b:EGFP-CAAX)* x *Tg(sox10:mRFP)*) that labels both neuronal membranes in the spinal cord (EGFP-CAAX) and oligodendrocyte membranes (mRFP). Using this line, we showed that the circular membrane features overlapped with the neuron cell membranes, suggesting that these circular membranes were indeed myelin that ectopically wrapped neuronal cell bodies (Fig. 6C).

Electron microscopy was then used to image selected regions in the spinal cord where neuronal cell bodies were more likely to occur in DomA-exposed fish (Fig. 6D). The DomA-exposed fish had neuronal cell bodies that were surrounded by additional membranes, which, together with the light microscopy data, could indicate ectopically myelinated cell bodies (Fig 6D2-D3).

### Pharmacologically reducing the number of oligodendrocytes results in lower numbers of circular myelin profiles

Findings from the genetic model (*kif1bp^-/-^*) suggest that oligodendrocytes wrap neuronal cell bodies when there is relatively less axonal surface area to myelinate.^59^ Thus, these circular profiles are said to result from the mismatch between the available myelinating oligodendrocytes and the axonal surface area. If the appearance of ectopically myelinated neuronal cell bodies was due to a higher number of oligodendrocytes relative to the axonal surface area, reducing the number of oligodendrocytes should also lead to the reduction in these ectopically myelinated neuronal cell bodies. To test this hypothesis, we used the small molecule GLI antagonist GANT61^58,59^ in (*Tg(mbp:EGFP)* x *Tg(mbp:EGFP-CAAX)*) fish to pharmacologically reduce the number of oligodendrocytes.

We first assessed whether GANT61 treatment successfully reduced the number of dorsal oligodendrocytes in the spinal cord. In control fish, exposure to all doses of GANT61 (10, 25, and 40 μM) led to significant reductions in oligodendrocyte number (Fig. 7A,E). For DomA-exposed fish, application of 25 and 40 μM GANT61 led to significant reductions in the oligodendrocyte numbers, while 10 μM did not (Fig. 7A,F).

**Figure 7:**
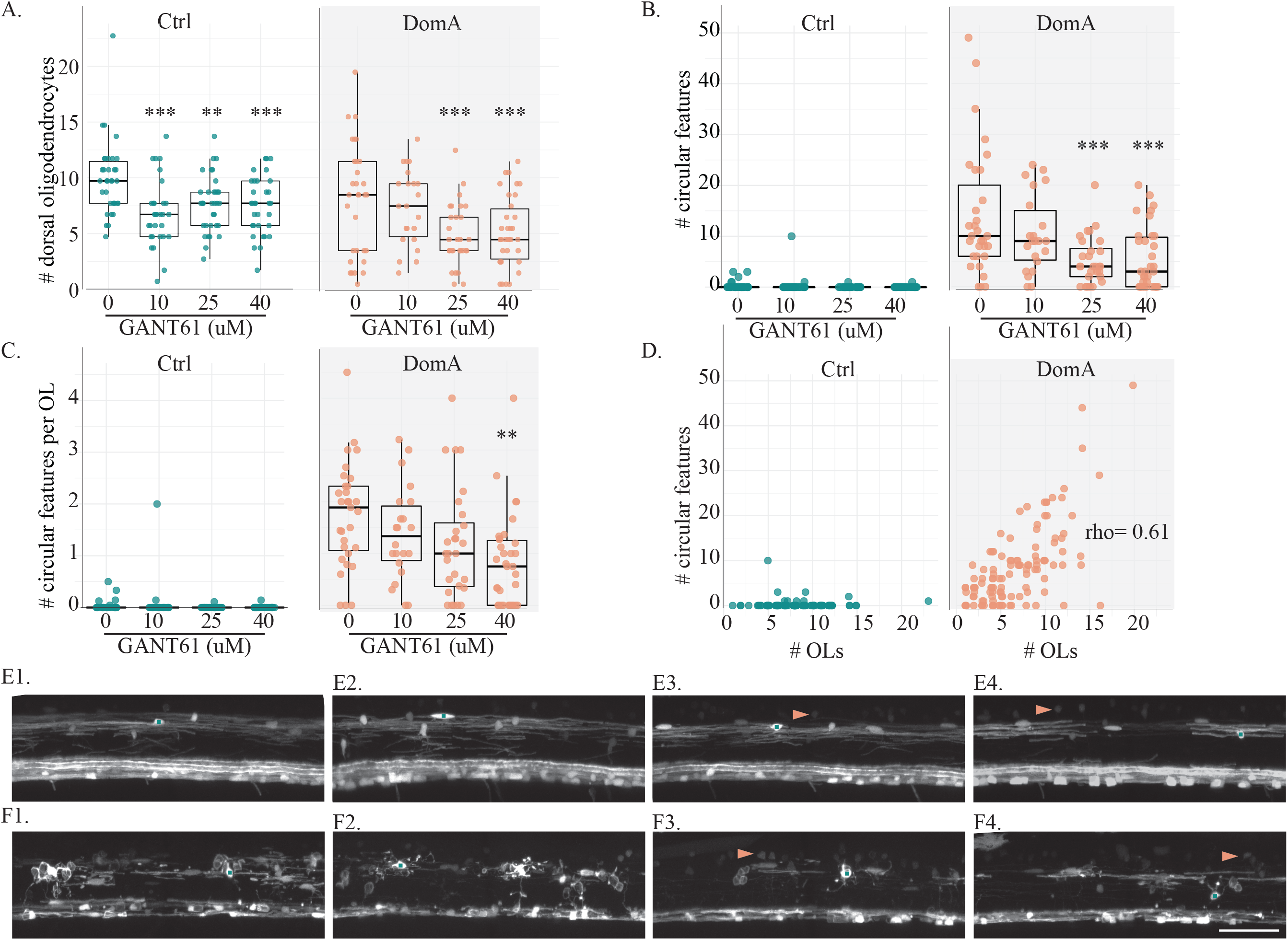
Treatment with GANT61, a small molecule that reduces the number of myelinating oligodendrocytes, also reduces the number of circular profiles in DomA treated fish. (A) *Tg(mbp:EGFP)* x *Tg(mbp:EGFP-CAAX)* double transgenic fish were exposed to DomA (0.14 ng) or vehicle at 2 dpf. Myelinating oligodendrocytes in the dorsal spinal cord were counted for control and DomA-exposed fish that were exposed to different concentrations of GANT61 (1.5-4 dpf). Each point represents the number of oligodendrocytes counted within a 269 μM imaging area in a single fish. (B) Number of odd circular features in dorsal spinal cord of the same control and DomA exposed fish plotted in Fig 7A. (C) The number of circular features per oligodendrocyte (OL) in the dorsal spinal cord of the control and DomA-exposed fish that were exposed to different concentrations of GANT61. (D) The number of circular features plotted against the number of dorsal OLs. (E) Images of control fish exposed to 0 μM (E1), 10 μM (E2), 25 μM (E3), and 40 μM GANT61 (E4). (F) Images of DomA fish exposed to 0 μM (F1), 10 μM (F2), 25 μM (F3), and 40 μM GANT61 (F4). Scale bar = 50 μm.* = p< 0.05, ** = p < 0.01, *** p < 0.001 by generalized mixed models with a Poisson or a negative binomial distribution. Data were aggregated from two trials. See Supplemental Fig. 4 for data from individual trials.

We then characterized the effects of GANT61 treatment on the number of putative ectopically myelinated neuronal cell bodies (Fig.7B,C). As expected, there were very few ectopically myelinated cell bodies in control fish, regardless of GANT61 treatment. In contrast, the majority of DomA-exposed fish had ectopically myelinated cell bodies. The number of ectopically-myelinated neuronal cell bodies was influenced by GANT61 treatment, with statistically significantly fewer ectopic features in DomA-exposed fish treated with 25 and 40 μM GANT61 compared to those that were not treated with GANT61 (25 μM GANT61–Estimate= −0.98, p= 0.0002; 40 μM GANT61–Estimate= −0.94, p= 0.0001) When taking into account the average number of ectopically myelinated cell bodies per oligogendrocyte, GANT61 treatment significantly reduced the number of circular features per oligodendrocytes at the highest dose (Fig 7C) (40 μM GANT61–Estimate = −0.786, p = 0.001).

Lastly, there was a slight positive correlation between the numbers of dorsal oligodendrocytes and the number of ectopically myelinated neuronal cell bodies (Fig. 7D) (rho = 0.611, p =3.96 e-13). The positive correlation was significant for both experimental trials. However, the first trial showed a stronger correlation between the number of oligodendrocytes and the ectopically myelinated neuronal cell bodies than the second one (Trial 1 – rho = 0.647, p = 4.06 e-08, Trial 2 – rho = 0.355, p = 0.007, Supplemental Fig. 4C, 4D).

## 4.5 DISCUSSION

The results reported here show that DomA exposure at 2 dpf leads to the loss of specific neuronal populations, including the Mauthner cells, prior to myelination (at 60 hpf). This indicates that the loss of these neurons is not secondary to myelin defects observed later in development (from 3.5 dpf onwards). Instead, the loss of reticulospinal neurons may contribute to the observed aberrant myelination pattern, characterized by both the overall reduction in myelin and the appearance of unusual circular myelin membranes.

Notably, DomA exposure at 2 dpf (but not at 1 dpf) led to the loss of Mauthner neurons. While Mauthner neurons are present at both 1 and 2 dpf, they have different intrinsic properties at different developmental stages, which in turn alter the neurons’ excitability and susceptibility to excitotoxicity. Thus, Mauthner cells are more excitable at 48 hpf versus 30 hpf, as measured by higher frequency and amplitude in their AMPA miniature excitatory postsynaptic potentials (mEPSPs).^61^ Furthermore, it has been suggested that these changes are due to a switch in AMPA receptor subunit composition between 30 hpf and 48 hpf. ^61^ It is conceivable that the developmental switch in AMPA receptor subunits of Mauthner neurons make these neurons more sensitive to DomA exposures later in development. It is well-known that DomA has different binding affinities for different AMPA and KA receptors that are composed of homomeric subunits.^7^ The switches in AMPA receptor subunits during development could change the sensitivity of neurons to DomA, making it more excitotoxic for the Mauthner neuron at the 2 dpf timepoint compared to the 1 dpf timepoint.

While DomA exposure at 2 dpf led to the loss of Mauthner cells at 2.5 dpf, it did not lead to any apparent losses in selected sensory neuron structures or the main axons of both the CaP and the RoP primary motor neurons. However, exposure to DomA did lead to significant losses in the MiP primary motor neurons. Primary motor neuron subtypes – CaP, RoP and MiP – have different axonal trajectories, innervate different regions of the muscle mass and have different electrical membrane properties.^62,63^ This may explain how DomA preferentially alters the main primary motor neuron axon of one subtype and not the others.

DomA exposure also affected the oligodendrocyte lineage. After the initial stages of myelination (2.5-3 dpf), DomA exposure led to significantly fewer myelinating oligodendrocytes in fish that also had severe myelin defects. DomA exposure also resulted in individual oligodendrocytes that produced fewer and shorter elongated myelin sheaths, while having more aberrant circular myelin membranes, which may be ectopically myelinated neuronal cell bodies (see below). In contrast, DomA did not alter the number of OPCs, which suggests that DomA does not alter OPC development. However, it is possible that DomA alters OPC developmental processes such as specification, proliferation, or survival in opposite ways that result in no changes to the total number of OPCs. Further experiments that directly investigate critical cellular processes including specification, migration, and differentiation would be necessary to confirm that DomA does not perturb OPC development.

The effects of DomA on the oligodendrocyte lineage may be secondary to the loss of reticulospinal neurons. Results from experiments using the *kif1bp* mutant^64^ were similar to those in this study. The loss of *kif1bp*, and the resulting reduction in axonal surface area, did not lead to fewer OPCs, nor did the axonal loss alter OPC specification, migration, or timing of differentiation. In contrast, the loss of *kif1bp* led to the impaired survival of oligodendrocytes and an overall reduction in myelination in the posterior region on the spinal cord where the axonal loss was more pronounced. This illustrates how target axons influence oligodendrocyte development, and conversely how the loss of axons leads to perturbations in oligodendrocyte survival, as well as oligodendrocyte loss.

The loss of the Mauthner axons may have contributed to the appearance of the unusual circular myelin membranes, which were tentatively identified as myelin that ectopically wrapped neuronal cell bodies. A recent study that used a genetic model that lacks reticulospinal axons (*kif1bp-/-*) reported an excess of ectopic myelin wrapping neuronal cell bodies.^59^ The authors posited that the oligodendrocytes myelinate neuronal cell bodies in cellular environments where there were fewer axons to myelinate relative to the number of oligodendrocytes present. To test this, they manipulated oligodendrocyte numbers using a pharmacological approach; decreasing the number of oligodendrocytes reduced the number of ectopically myelinated neuronal cell bodies while increasing the number of oligodendrocytes increased the number of these ectopic features. We hypothesized that DomA-induced axonal loss led to these circular myelin membranes by a similar mechanism. Treatment of DomA-exposed fish with GANT61 – a small molecule that decreases oligodendrocyte number – caused a reduction in the number of oligodendrocytes as well as the circular myelin membranes. This supports the hypothesis that these circular myelin membranes arise in DomA-treated fish due to the loss of reticulospinal axons and the resulting mismatch of axonal surface area to oligodendrocyte number, which causes oligodendrocytes to ectopically myelinate neuronal cell bodies.

The results of our study point to reticulospinal neurons as targets of DomA. We cannot rule out the possibility that DomA also targets the oligodendrocyte lineage directly. It is possible that DomA directly binds to ionotrophic glutamate receptors in oligodendrocytes, potentially inhibiting their ability to myelinate axons and leading to further axonal loss. Nevertheless, our results strongly support the idea that reticulospinal neurons are the initial targets of DomA and that their loss contributes to the myelination defects observed.

## 4.6 CONCLUSION

This study identified neuronal subpopulations (Mauthner cell, MiP primary motor neurons) as the initial targets for DomA exposure that occurs during the developmental period prior to myelination (proposed model, Fig. 8). The loss of these neurons, and the resulting changes to the axonal environment in the spinal cord, contributed to the pronounced myelin defects observed. DomA also led to loss of oligodendrocytes and the inability of remaining oligodendrocytes to appropriately myelinate targets in the spinal cord. This may be a secondary effect due to the initial loss of neurons or to direct effects of DomA on oligodendrocytes. These results suggest that DomA exposure can target specific neuronal populations and alter cellular environments that lead to pronounced and lasting structural and behavioral phenotypes.

**Figure 8:**
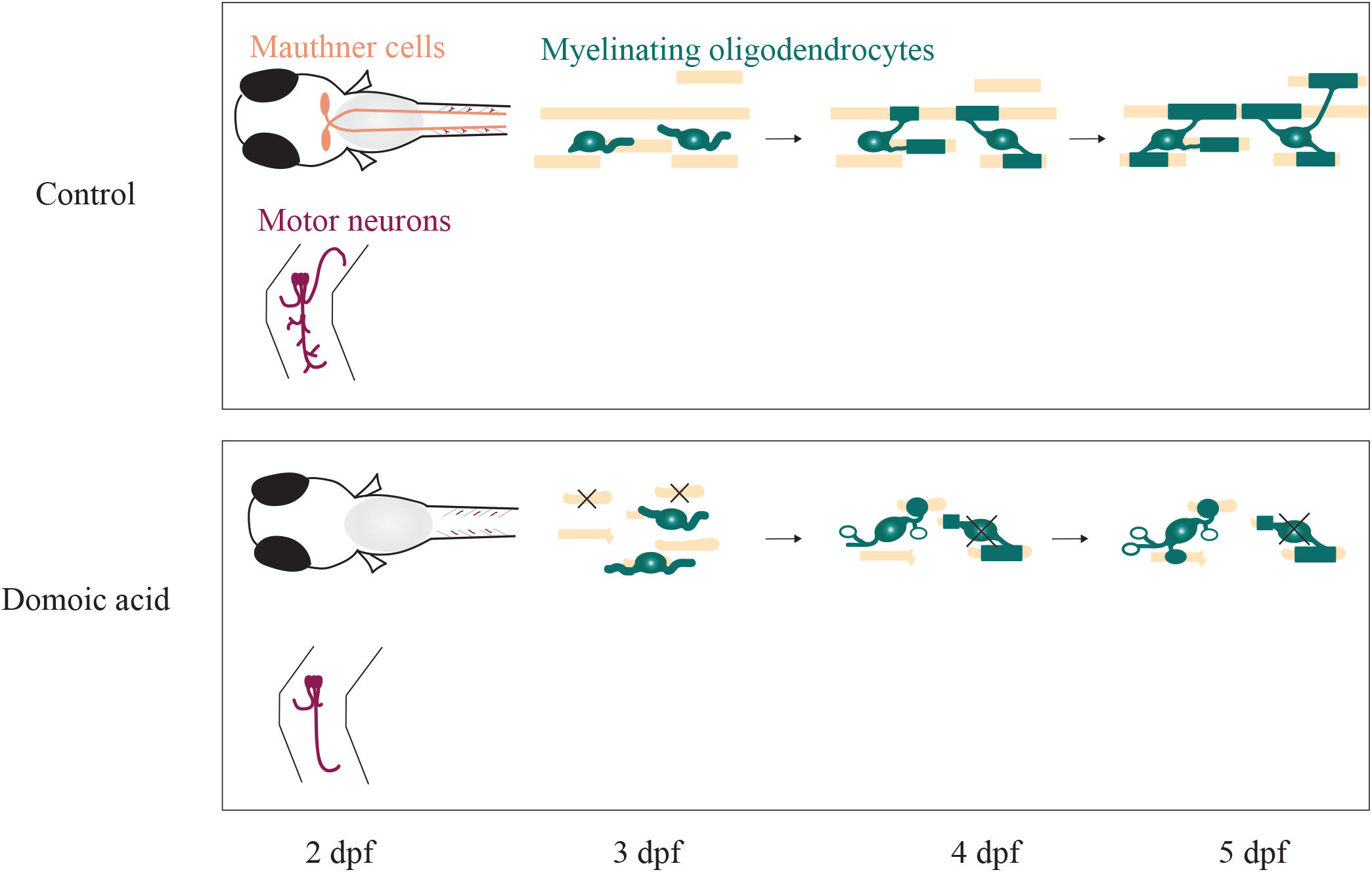
Proposed model. A. During normal development, the Mauthner cells are present at 2.5 dpf, and axon wrapping and myelination commences at 2.5 dpf. By 5 dpf, nascent myelin sheath formation is completed, and robust startle behavior can be observed. B. DomA exposure results in the loss of Mauthner cells and selected primary motor neuron subtypes. This results in the reduction in axonal surface area in the spinal cord prior to myelination. With the reduced axonal surface area, a subset of cells from the oligodendrocyte lineage inappropriately myelinate neuronal cell bodies while others undergo cell death. The loss of the Mauthner cells, the reduction in primary motor neuron collaterals, and the myelination defects could all potentially contribute to the startle deficits observed in the larval stages.

## ACKNOWLEDGEMENTS

We would like to thank Dr. Matthew Salanga (Northern Arizona University) for training and advice regarding plasmid design and transgenic line generation, Dr. Louis Kerr (MBL microscopy facility) and Dr. Nadja Brun for light microscopy training and advice, Dr. Eduardo Rosa-Molinar, Dr. Noraida Marinez-Rivera, and Irma Torres-Vazquez for electron microscopy training and advice, and several labs for generously providing us with zebrafish transgenic lines and plasmid constructions that made this work possible: Dr. Sarah Kucenas (University of Virginia), Dr. Bruce Appel (University of Colorado, Denver), Dr. Kelly Monk (Vollum Institute, Oregon Health and Sciences University), Dr. Charles Kaufman (University of Washington St. Louis), Dr. Leonard Zon (Harvard Medical School), Dr. David Lyons (University of Edinburg), and Dr. David Grunwald (University of Utah).

## AUTHOR CONTRIBUTIONS

JMP wrote the manuscript, designed the experiments, generated the data, and performed all the analyses. KMH prepped and imaged the samples for the electron microscopy. NA and MEH obtained the funding and assisted in the experimental design. All authors contributed to editing the manuscript. All authors agree to accountability for the content.

## DATA AVAILABILITY

The data that support the current study are available from the corresponding author upon reasonable request.

## REFERENCES

1. Hoagland, P., Anderson, D. M., Kaoru, Y. & White, A. W. The economic effects of harmful algal blooms in the United States: Estimates, assessment issues, and information needs. Estuaries 25, 819–837 (2002).

2. Anderson, D. M. et al. Marine harmful algal blooms (HABs) in the United States: History, current status and future trends. Harmful Algae 102, (2021).

3. Berdalet, E. et al. Marine harmful algal blooms, human health and wellbeing: challenges and opportunities in the 21st century. J. Mar. Biol. Assoc. United Kingdom 96, 61–91 (2016).

4. Grattan, L. M., Holobaugh, S. & Morris, J. G. Harmful algal blooms and public health. Harmful Algae 57, 2–8 (2016).

5. Backer, L., LE, F., AD, R. & DG, B. Epidemiology, public health and human diseases associated with harmful marine algae. in Manual on Harmful Marine Microalgae (eds. GM, H., DM, A. & AD, C.) 723–749 (UNESCO Publishing, 2003).

6. Stewart, G. R., Zorumski, C. F., Price, M. T. & Olney, J. W. Domoic acid: A dementia-inducing excitotoxic food poison with kainic acid receptor specificity. Exp. Neurol. 110, 127–138 (1990).

7. Hampson, D. R. & Manalo, J. L. The activation of glutamate receptors by kainic acid and domoic acid. Nat. Toxins 6, 153–8 (1998).

8. Bates, S. S. et al. Pennate Diatom *Nitzschia pungens* as the Primary Source of Domoic Acid, a Toxin in Shellfish from Eastern Prince Edward Island, Canada. Can. J. Fish. Aquat. Sci. 46, 1203–1215 (1989).

9. Fritz, L., Quilliam, M. A., Wright, J. L. C., Beale, A. M. & Work, T. M. An outbreak of domoic acid poisoning attributed to the pennate diatom *Pseudonitzchia australis*. J. Phycol. 28, 439–442 (1992).

10. Nakajima, S. & Potvin, J. L. Neural and behavioural effects of domoic acid, an amnesic shellfish toxin, in the rat. Can. J. Psychol. 46, 569–81 (1992).

11. Perl, T. M. et al. An outbreak of toxic encephalopathy caused by eating mussels contaminated with domoic acid. N. Engl. J. Med. 322, 1775–80 (1990).

12. Petroff, R. et al. Public health risks associated with chronic, low-level domoic acid exposure: A review of the evidence. Pharmacol. Ther. 227, (2021).

13. Wekell, J. C., Jurst, J. & Lefebvre, K. A. The origin of the regulatory limits for PSP and ASP toxins in shellfish. J. Shellfish Res. 23, 927–930 (2010).

14. Mariën, K. Establishing tolerable dungeness crab (Cancer magister) and razor clam (Siliqua patula) domoic acid contaminant levels. Environ. Health Perspect. 104, 1230–6 (1996).

15. Bernard, P. B., MacDonald, D. S., Gill, D. A., Ryan, C. L. & Tasker, R. A. Hippocampal mossy fiber sprouting and elevated trkB receptor expression following systemic administration of low dose domoic acid during neonatal development. Hippocampus 17, 1121–1133 (2007).

16. Perry, M. A., Ryan, C. L. & Tasker, R. A. Effects of low dose neonatal domoic acid administration on behavioural and physiological response to mild stress in adult rats. Physiol. Behav. 98, 53–59 (2009).

17. Gill, D. A. et al. Neonatal exposure to low-dose domoic acid lowers seizure threshold in adult rats. Neuroscience 169, 1789–1799 (2010).

18. Mills, B. D. et al. Prenatal domoic acid exposure disrupts mouse pro-social behavior and functional connectivity MRI. Behav. Brain Res. 308, 14–23 (2016).

19. Shiotani, M. et al. Neurobehavioral assessment of mice following repeated oral exposures to domoic acid during prenatal development. Neurotoxicol. Teratol. 64, 8–19 (2017).

20. Adams, A. L., Doucette, T. A., James, R. & Ryan, C. L. Persistent changes in learning and memory in rats following neonatal treatment with domoic acid. Physiol. Behav. 96, 505–12 (2009).

21. Doucette, T. A., Bernard, P. B., Yuill, P. C., Tasker, R. A. & Ryan, C. L. Low doses of non-NMDA glutamate receptor agonists alter neurobehavioural development in the rat. Neurotoxicol. Teratol. 25, 473–479 (2003).

22. Burt, M. A., Ryan, C. L. & Doucette, T. A. Altered responses to novelty and drug reinforcement in adult rats treated neonatally with domoic acid. Physiol. Behav. 93, 327–336 (2008).

23. Tasker, R. A. R., Perry, M. A., Doucette, T. A. & Ryan, C. L. NMDA receptor involvement in the effects of low dose domoic acid in neonatal rats. Amino Acids 28, 193–196 (2005).

24. Ryan, C. L. et al. Altered social interaction in adult rats following neonatal treatment with domoic acid. Physiol. Behav. 102, 291–295 (2011).

25. Marriott, A. L., Ryan, C. L. & Doucette, T. A. Neonatal domoic acid treatment produces alterations to prepulse inhibition and latent inhibition in adult rats. Pharmacol. Biochem. Behav. 103, 338–344 (2012).

26. Burt, M. A., Ryan, C. L. & Doucette, T. A. Low dose domoic acid in neonatal rats abolishes nicotine induced conditioned place preference during late adolescence. Amino Acids 35, 247–249 (2008).

27. Panlilio, J. M., Jones, I. T., Salanga, M. C., Aluru, N. & Hahn, M. E. Developmental Exposure to Domoic Acid Disrupts Startle Response Behavior and Circuitry in Zebrafish. Toxicol. Sci. 182, 310–326 (2021).

28. Panlilio, J. M., Aluru, N. & Hahn, M. E. Developmental Neurotoxicity of the Harmful Algal Bloom Toxin Domoic Acid: Cellular and Molecular Mechanisms Underlying Altered Behavior in the Zebrafish Model. Environ. Health Perspect. 128, 117002 (2020).

29. Nave, K.-A. & Trapp, B. D. Axon-Glial Signaling and the Glial Support of Axon Function. Annu. Rev. Neurosci. 31, 535–561 (2008).

30. Nave, K.-A. Myelination and support of axonal integrity by glia. Nature 468, 244–52 (2010).

31. Bradl, M. & Lassmann, H. Oligodendrocytes: biology and pathology. Acta Neuropathol. 119, 37–53 (2010).

32. Park, H.-C., Shin, J., Roberts, R. K. & Appel, B. An olig2 reporter gene marks oligodendrocyte precursors in the postembryonic spinal cord of zebrafish. Dev. Dyn. 236, 3402–7 (2007).

33. Patneau, D. K., Wright, P. W., Winters, C., Mayer, M. L. & Gallo, V. Glial cells of the oligodendrocyte lineage express both kainate-and AMPA-preferring subtypes of glutamate receptor. Neuron 12, 357–371 (1994).

34. Bergles, D. E., Roberts, J. D., Somogyi, P. & Jahr, C. E. Glutamatergic synapses on oligodendrocyte precursor cells in the hippocampus. Nature 405, 187–91 (2000).

35. Yuan, X. et al. A role for glutamate and its receptors in the regulation of oligodendrocyte development in cerebellar tissue slices. Development 125, 2901–14 (1998).

36. Barres, B. A., Koroshetz, W. J., Swartz, K. J., Chun, L. L. Y. & Corey, D. P. Ion channel expression by white matter glia: the O-2A glial progenitor cell. Neuron 4, 507–524 (1990).

37. Borges, K., Ohlemeyer, C., Trotter, J. & Kettenmann, H. AMPA/kainate receptor activation in murine oligodendrocyte precursor cells leads to activation of a cation conductance, calcium influx and blockade of delayed rectifying K+ channels. Neuroscience 63, 135–149 (1994).

38. Alberdi, E., Sánchez-Gómez, M. V., Marino, A. & Matute, C. Ca(2+) influx through AMPA or kainate receptors alone is sufficient to initiate excitotoxicity in cultured oligodendrocytes. Neurobiol. Dis. 9, 234–43 (2002).

39. Matute, C., Domercq, M. & Sánchez-Gómez, M.-V. Glutamate-mediated glial injury: mechanisms and clinical importance. Glia 53, 212–24 (2006).

40. Rosenberg, P. A. et al. Mature myelin basic protein-expressing oligodendrocytes are insensitive to kainate toxicity. J. Neurosci. Res. 71, 237–45 (2003).

41. Deng, W., Rosenberg, P. a, Volpe, J. J. & Jensen, F. E. Calcium-permeable AMPA/kainate receptors mediate toxicity and preconditioning by oxygen-glucose deprivation in oligodendrocyte precursors. Proc. Natl. Acad. Sci. U. S. A. 100, 6801–6 (2003).

42. Gallo, V. et al. Oligodendrocyte progenitor cell proliferation and lineage progression are regulated by glutamate receptor-mediated K+ channel block. J. Neurosci. 16, 2659–2670 (1996).

43. Matute, C. Characteristics of acute and chronic kainate excitotoxic damage to the optic nerve. Proc. Natl. Acad. Sci. U. S. A. 95, 10229–34 (1998).

44. Almeida, R. G., Czopka, T., Ffrench-Constant, C. & Lyons, D. A. Individual axons regulate the myelinating potential of single oligodendrocytes in vivo. Development 138, 4443–50 (2011).

45. Kucenas, S. et al. CNS-derived glia ensheath peripheral nerves and mediate motor root development. Nat. Neurosci. 11, 143–151 (2008).

46. Kucenas, S., Snell, H. & Appel, B. nkx2.2a promotes specification and differentiation of a myelinating subset of oligodendrocyte lineage cells in zebrafish. Neuron Glia Biol. 4, 71–81 (2008).

47. Takada, N., Kucenas, S. & Appel, B. Sox10 is necessary for oligodendrocyte survival following axon wrapping. Glia 58, 996–1006 (2010).

48. White, R. M. et al. DHODH modulates transcriptional elongation in the neural crest and melanoma. Nature 471, 518 (2011).

49. Cianciolo Cosentino, C., Roman, B. L., Drummond, I. A. & Hukriede, N. A. Intravenous microinjections of zebrafish larvae to study acute kidney injury. J. Vis. Exp. (2010). doi:10.3791/2079

50. Ripley, B. et al. Package ‘MASS’. (2018).

51. Jung, S.-H. et al. Visualization of myelination in GFP-transgenic zebrafish. Dev. Dyn. 239, 592–597 (2010).

52. Kwan, K. M. et al. The Tol2kit: A multisite gateway-based construction kit forTol2 transposon transgenesis constructs. Dev. Dyn. 236, 3088–3099 (2007).

53. Hoshijima, K., Jurynec, M. J. & Grunwald, D. J. Precise Editing of the Zebrafish Genome Made Simple and Efficient. Dev. Cell 36, 654–67 (2016).

54. Longair, M. H., Baker, D. A. & Armstrong, J. D. Simple Neurite Tracer: open source software for reconstruction, visualization and analysis of neuronal processes. Bioinformatics 27, 2453–2454 (2011).

55. Wobbrock, J. O., Findlater, L., Gergle, D. & Higgins, J. J. The aligned rank transform for nonparametric factorial analyses using only anova procedures. in Proceedings of the 2011 annual conference on Human factors in computing systems - CHI 2011 143 (ACM Press, 2011). doi:10.1145/1978942.1978963

56. Karlsson, J., von Hofsten, J. & Olsson, P.-E. Generating Transparent Zebrafish: A Refined Method to Improve Detection of Gene Expression During Embryonic Development. Mar. Biotechnol. 3, 0522–0527 (2001).

57. Inoue, D. & Wittbrodt, J. One for all--a highly efficient and versatile method for fluorescent immunostaining in fish embryos. PLoS One 6, e19713 (2011).

58. Agyeman, A., Jha, B. K., Mazumdar, T. & Houghton, J. A. Mode and specificity of binding of the small molecule GANT61 to GLI determines inhibition of GLI-DNA binding. Oncotarget 5, 4492–503 (2014).

59. Almeida, R. G. et al. Myelination of Neuronal Cell Bodies when Myelin Supply Exceeds Axonal Demand. Curr. Biol. 28, 1296–1305.e5 (2018).

60. Hothorn, T., Bretz, F. & Westfall, P. Simultaneous Inference in General Parametric Models. Biometrical J. 50, 346–363 (2007).

61. Patten, S. A. & Ali, D. W. AMPA receptors associated with zebrafish Mauthner cells switch subunits during development. J. Physiol. 581, 1043–56 (2007).

62. Babin, P. J., Goizet, C. & Raldúa, D. Zebrafish models of human motor neuron diseases: Advantages and limitations. Prog. Neurobiol. 118, 36–58 (2014).

63. Moreno, R. L. & Ribera, A. B. Zebrafish motor neuron subtypes differ electrically prior to axonal outgrowth. J. Neurophysiol. 102, 2477–84 (2009).

64. Almeida, R. & Lyons, D. Oligodendrocyte Development in the Absence of Their Target Axons In Vivo. PLoS One 11, e0164432 (2016).

