## Supplemental_information for "Developmental exposure to domoic acid targets reticulospinal neurons and leads to aberrant myelination in the spinal cord"

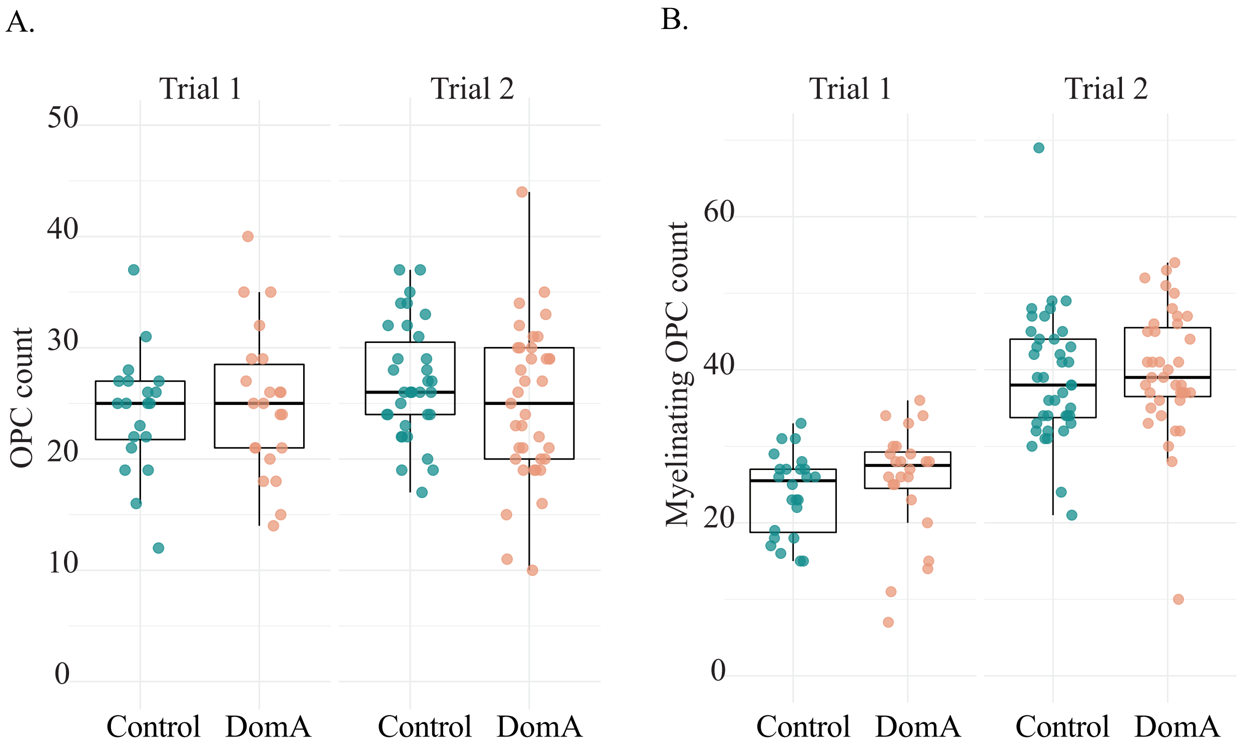


**Supplemental Figure 1: Trial differences in oligodendrocyte precursor cell counts.**

**Supplemental Figure 1: Trial differences in oligodendrocyte precursor cell counts**

(A) Oligodendrocyte precursor cell (OPC) count in the dorsal spinal cords in the double transgenic fish, *Tg(sox10:mRFP)* x *Tg(olig2:EGFP),* imaged at 2.5 dpf. Each point represents the number of OPCs counted within the 403.1µM imaging area in a single fish. There were no differences in OPC count based on treatment or on repeat trials.

(B) Oligodendrocyte precursor cell (OPC) count in dorsal spinal cords of the 354.3µM long region in the dorsal spinal cords in the double transgenic fish, *Tg(sox10:RFP)* x *Tg(nkx2.2a:mEGFP),* imaged at 2.5 dpf. Each point represents the number of OPCs counted within the 354.3µM imaging area in a single fish. While there were no differences in OPC count based on treatment, there were differences in OPC count between the two repeat trials.

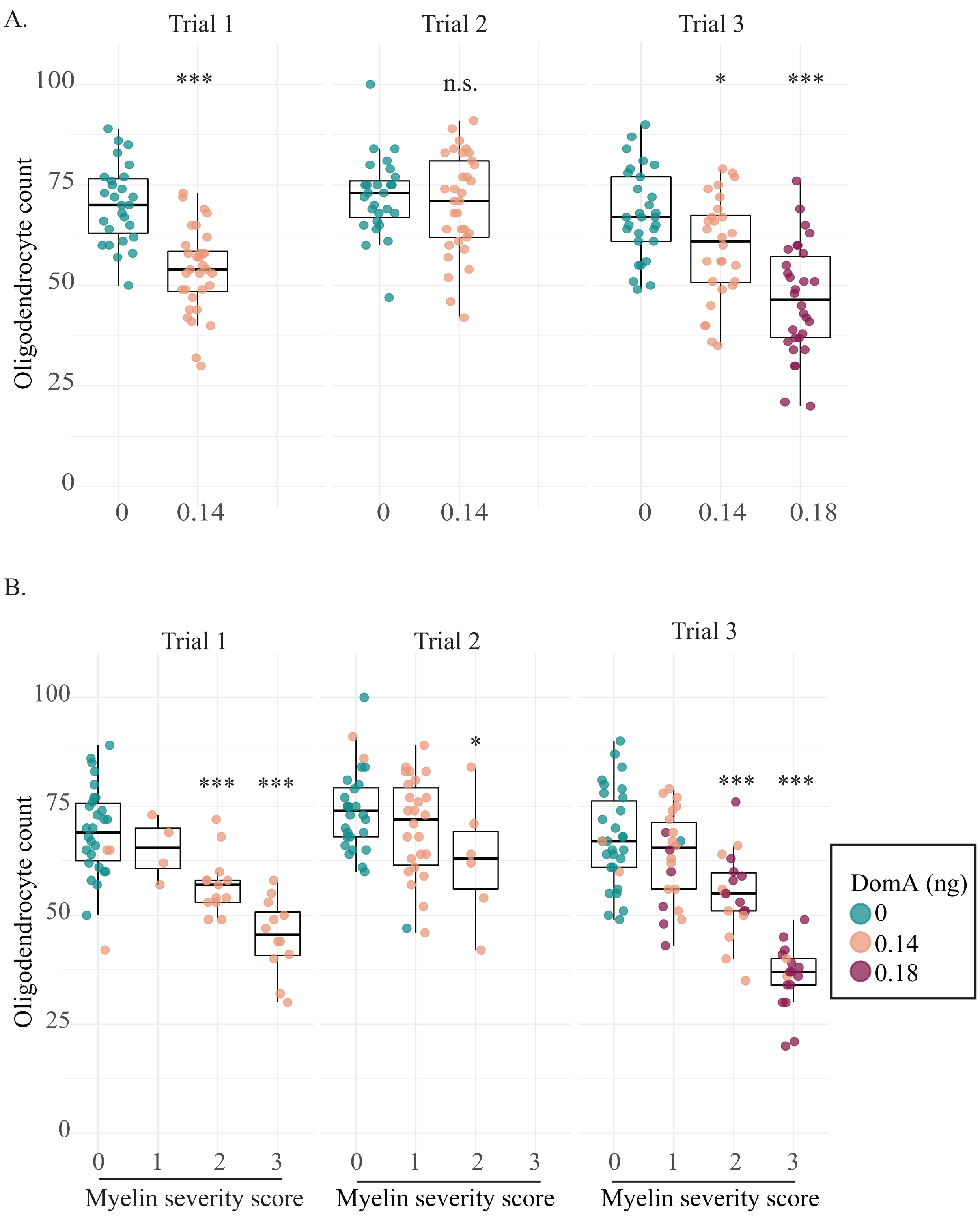


**Supplemental Figure 2: Trial differences in myelinating oligodendrocyte cell counts**

**Supplemental Figure 2: Trial differences in myelinating oligodendrocyte cell counts**

(A) Myelinating oligodendrocyte count in the spinal cords. *Tg(mbp:EGFP)* imaged at 2.5 dpf. Individual points represent myelinating oligodendrocyte counts within the 403.9µM imaging area in a single fish. Trial 2 shows no significant differences between control (0 ng DomA) and domoic acid exposed fish (0.14 ng DomA). In 2 out of the 3 trials, DomA treatment significantly reduced the number of oligodendrocytes (DomA-exposed larvae (0.14 ng) (Coefficient = -0.268, Error= 0.043, p = 6.13 e-10 and Coefficient = -0.136, Error = 0.059, p=0.02 respectively).

Fish were further subdivided by the severity of the myelin defect observed. No treated fish in Trial 2 had myelin with the most severe score (category 3)

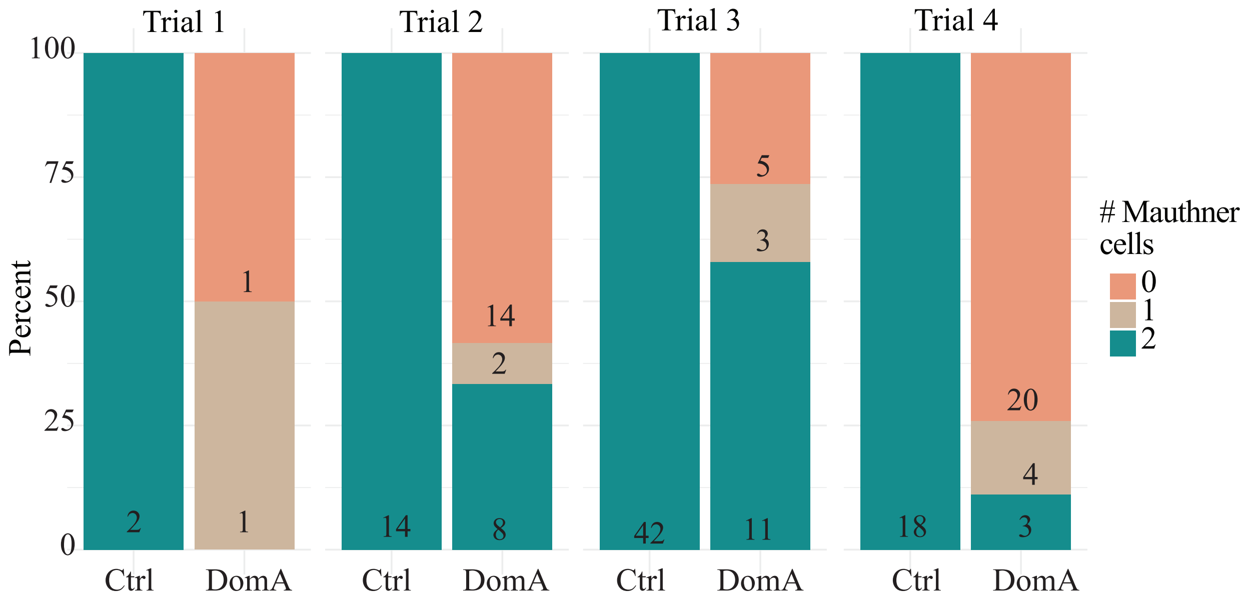


**Supplemental Figure 3: Trial differences in Mauthner cell counts.**

**Supplemental Figure 3: Trial differences in Mauthner cell counts.**
Percentage of embryos exposed to DomA (2 dpf) or vehicle that had 0, 1, or 2 Mauthner cells as visualized by staining with anti-3A10 at 60 hpf. Numbers of larvae with each phenotype are listed within each bar. Data for the individual experimental trials are shown.The degree of DomA-induced losses in Mauthner cells varied among repeated experiments. In 3 out of 4 trials, the majority of DomA-exposed fish had 0 out of the 2 Mauthner cells (Supplemental Fig. 3). Nonetheless, iIn all trials, DomA-exposed fish showed Mauthner cell numbers that were significantly different from controls ($p < 10^{-13}$).

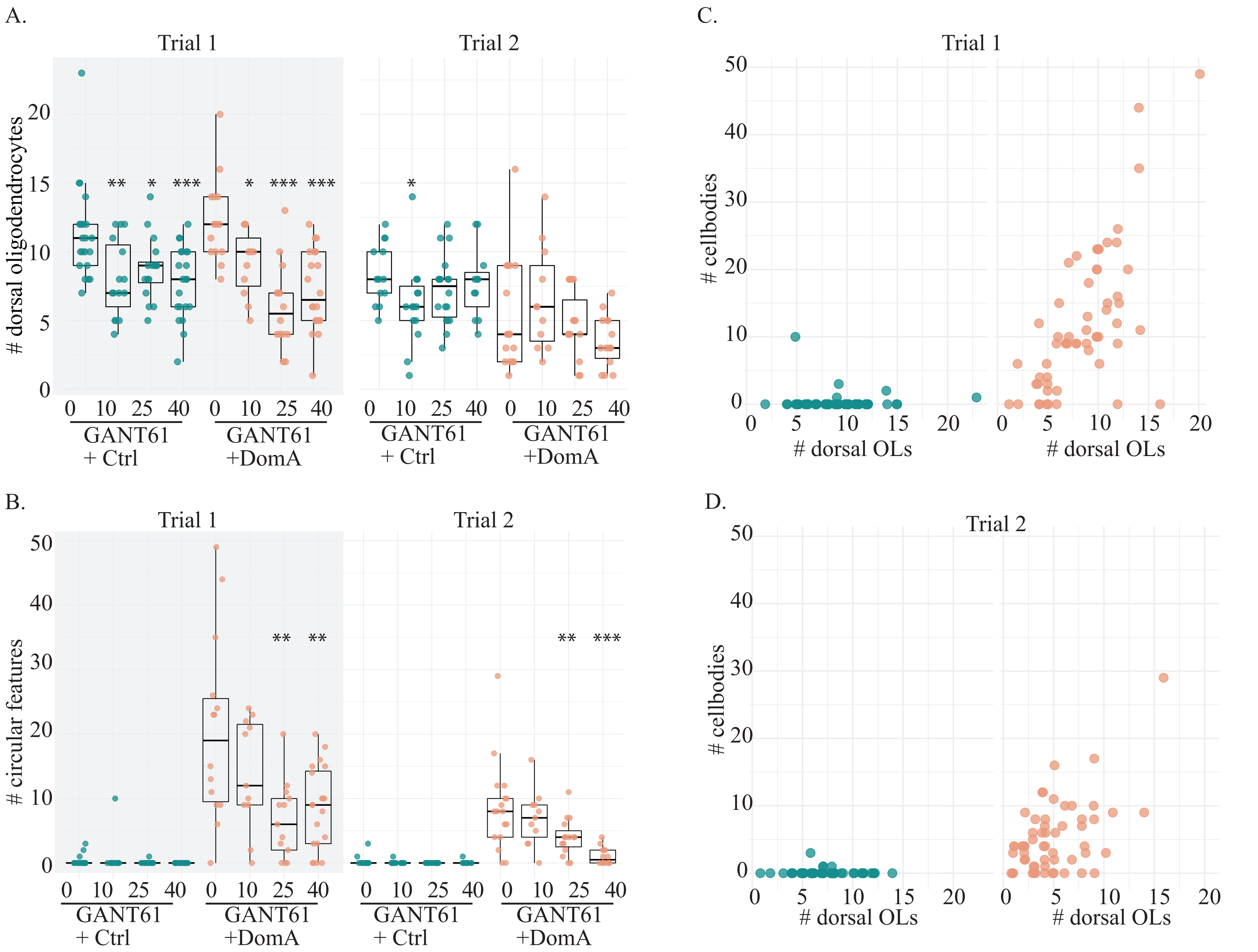


**Supplemental Figure 4: Trial differences in GANT61 treatment**

**Supplemental Figure 4: Trial differences in GANT61 treatment**

(A) Number of myelinating oligodendrocytes in dorsal spinal cord of control and DomA-exposed fish that were also exposed to different concentrations of GANT61 (1.5- 4 dpf). Each point represents the number of oligodendrocytes counted within a 269 μM imaging area in a single fish.

(B) Total number of circular features in dorsal spinal cord of the control and DomA-exposed fish that were exposed to different concentrations of GANT61.

(C) The number of circular features plotted against the number of dorsal oligodendrocytes in Trial 1

(D) The number of circular features plotted against the number of dorsal oligodendrocytes in Trial 2

* = $p < 0.05$, ** = $p < 0.01$, *** $p < 0.001$ by generalized mixed models with a Poisson or a negative binomial distribution.
